# T cell receptor is required for differentiation but not maintenance of intestinal intraepithelial lymphocytes

**DOI:** 10.1101/2020.06.04.134510

**Authors:** Angelina M. Bilate, Mariya London, Tiago B. R. Castro, Luka Mesin, Suppawat Kongthong, Audrey Harnagel, Gabriel D. Victora, Daniel Mucida

**Affiliations:** Laboratory of Mucosal Immunology, The Rockefeller University, New York, NY 10065, USA; Laboratory of Lymphocyte Dynamics, The Rockefeller University, New York, NY 10065, USA

## Abstract

The gut epithelium is populated by intraepithelial lymphocytes (IELs), a heterogeneous T cell population with cytotoxic and regulatory properties. Migrating peripheral CD4^+^ T cells, including regulatory (Treg) and conventional T cells (Tconv), acquire an IEL (CD4-IEL) program upon arrival at the epithelium. However, the specific role of the T cell receptor (TCR) in this process remains unclear. Single-cell TCR repertoire and transcriptomic analysis of intraepithelial CD4^+^ T cells revealed different extents of clonal expansion and TCR overlap between cell states; fully differentiated CD4-IELs from Tregs or Tconvs were the least diverse. Conditional deletion of TCR on differentiating CD4^+^ T cells or of MHCII on intestinal epithelial cells prevented CD4-IEL differentiation. However, TCR ablation on developed CD4-IELs did not affect their accumulation. These results indicate that local recognition of a limited set of antigens is an essential signal for the differentiation and adaptation of T cells to the epithelium.

## Introduction

T cell receptor (TCR) diversity of circulating T lymphocytes is ontogenically determined by thymic selection, but T cell localization and repertoire are largely dependent on exposure to, and recognition of, tissue-specific antigens (Hogquist and Jameson, 2014). In addition to these features, tissue imprinting influences T cell subset differentiation and function (Faria et al., 2017). In the intestine, T cells located in the lamina propria (LP), and in the intraepithelial (IE) compartment display gut-specific characteristics, such as expression of gut-homing integrins, but differ markedly in TCR usage, migration patterns and function (McDonald et al., 2018).

Intestinal intraepithelial lymphocytes (IELs) comprise a heterogeneous population of T cells, yet they all share common features such as tissue residency, cytotoxic potential, activated phenotype, and expression of CD8αα homodimers (Cheroutre et al., 2011; McDonald et al., 2018). Previous studies propose that CD8αα homodimers, in contrast to conventional CD8αβ heterodimers, work as TCR co-repressors by binding to thymus leukemia (TL) antigen expressed on epithelial cells, resulting in decreased antigen sensitivity of the TCR (Cheroutre and Lambolez, 2008). Similar to regulatory T cells (Treg) and invariant natural killer cells (iNKT), natural IELs (nIEL) such as CD8αα^+^TCRαβ^+^ are agonist-selected in the thymus (Leishman et al., 2002; Moran et al., 2011; Yamagata et al., 2004), suggesting an important role for TCR specificity in IEL development. Additionally, transgenic mice carrying an αβTCR from naturally-occurring CD8αα^+^TCRαβ^+^ T cells preferentially differentiate towards this lineage, further indicating that TCR signaling strength itself may drive IEL fate (Mayans et al., 2014; McDonald et al., 2015; McDonald et al., 2014).

In addition to the developmentally-imprinted TCR features of intestinal T cells, luminal stimulation and other gut-enriched factors, such as microbial metabolites and epithelial cell factors, further influence peripheral T cell differentiation and plasticity (Atarashi et al., 2013; Lathrop et al., 2011; Sujino et al., 2016; Xu et al., 2018; Yang et al., 2014). Conventional CD4^+^ T cells and Tregs can differentiate into CD4-IELs expressing CD8αα in a microbiota-dependent manner upon migration to the gut epithelium (Bilate et al., 2016; Sujino et al., 2016). However, the role of TCR signaling in IEL differentiation, location and function has not been established. This is of particular interest given the relatively low abundance of MHC class II-expressing cells in the gut epithelium and other barrier surfaces (Faria et al., 2017).

We sought to address how TCR properties and signaling modulate the location and plasticity of CD4^+^ T cells in the intestine. We combined TCR repertoire analysis with single-cell transcriptomics using a fate-mapping strategy that allowed us to track Tregs and conventional T cells (Tconv) as they migrate to the intestinal epithelium and differentiate into CD8αα-expressing IELs. Reduced TCR diversity was associated with terminal differentiation of CD4^+^ T cells into CD4-IELs; Tregs and intermediate stages were more diverse, whereas fully differentiated CD4-IELs were clonally restricted. Using *in vivo* genetic tools, we showed that ablation of surface TCR complexes on Tregs and other activated CD4^+^ T cells impaired CD4-IEL differentiation, suggesting that TCR expression is required for terminal T cell differentiation at the intestinal epithelium. Inducible deletion of MHC class II on intestinal epithelial cells (IEC) also prevented CD4-IEL differentiation, resulting in Treg accumulation at the epithelium. However, TCR ablation in fully differentiated CD4-IELs had little, if any, impact on their accumulation or on the maintenance of the IEL transcriptional program. Our findings indicate that TCR expression and local MHC class II on IECs are required for T cell plasticity at the intestinal epithelium, but not for the maintenance of the IEL program.

## Results

### Clonal expansion of intraepithelial CD4^+^ T cells

Specific TCR usage has been reported in both natural- and peripherally-induced Tregs located in distinct tissues, including the intestine (Fan and Rudensky, 2016; Lathrop et al., 2011; Liu et al., 2009; Zhou et al., 2009). While in most sites Tregs are thought to stably express Foxp3, we previously showed that a fraction of Tregs loses Foxp3 and acquires an IEL program, including CD8αα expression, upon migrating to the gut epithelium (Sujino et al., 2016). To define to what extent such plasticity is associated with specific TCR usage, we first analyzed the diversity of CD4^+^ T cells in the small intestine epithelium in wild-type (WT) mice using inducible Foxp3 fate-mapping coupled with Foxp3-reporter approaches. By crossing *Foxp3*^*eGFP-*CreERT2^ *Rosa26*^|*s*|-td-Tomato^ (i*Foxp3*^Tom^) with *Foxp3*^GFP^ mice (i*Foxp3*^Tom^ *Foxp3*^GFP^), we were able to specifically analyze the TCRαβ repertoire of four distinct subsets of IELs upon continuous tamoxifen administration: conventional CD4^+^ T cells (CD8αα^−^, Tomato^−^, GFP^−^; Tconv), current Tregs (CD8αα^−^, GFP^+^) and CD4-IELs originated from either Tconv (CD8αα^+^, Tomato^−^) or from Tregs (CD8αα^+^, Tomato^+^; ex-Treg CD4-IEL) (Figure S1A). We compared the IEL TCR repertoire diversity with Tregs isolated from gut-draining mesenteric lymph nodes (mLN) and lamina propria from the same animals by single-cell TCRαβ sequencing (scTCRseq; Figure 1A). Cells with identical TCRβ CDR3 nucleotide sequences were considered as the same clones; additionally, clonality based on TCRβ CDR3s was confirmed by sequencing TCRα of the same expanded cells (Figure S1B).

**Figure 1.**
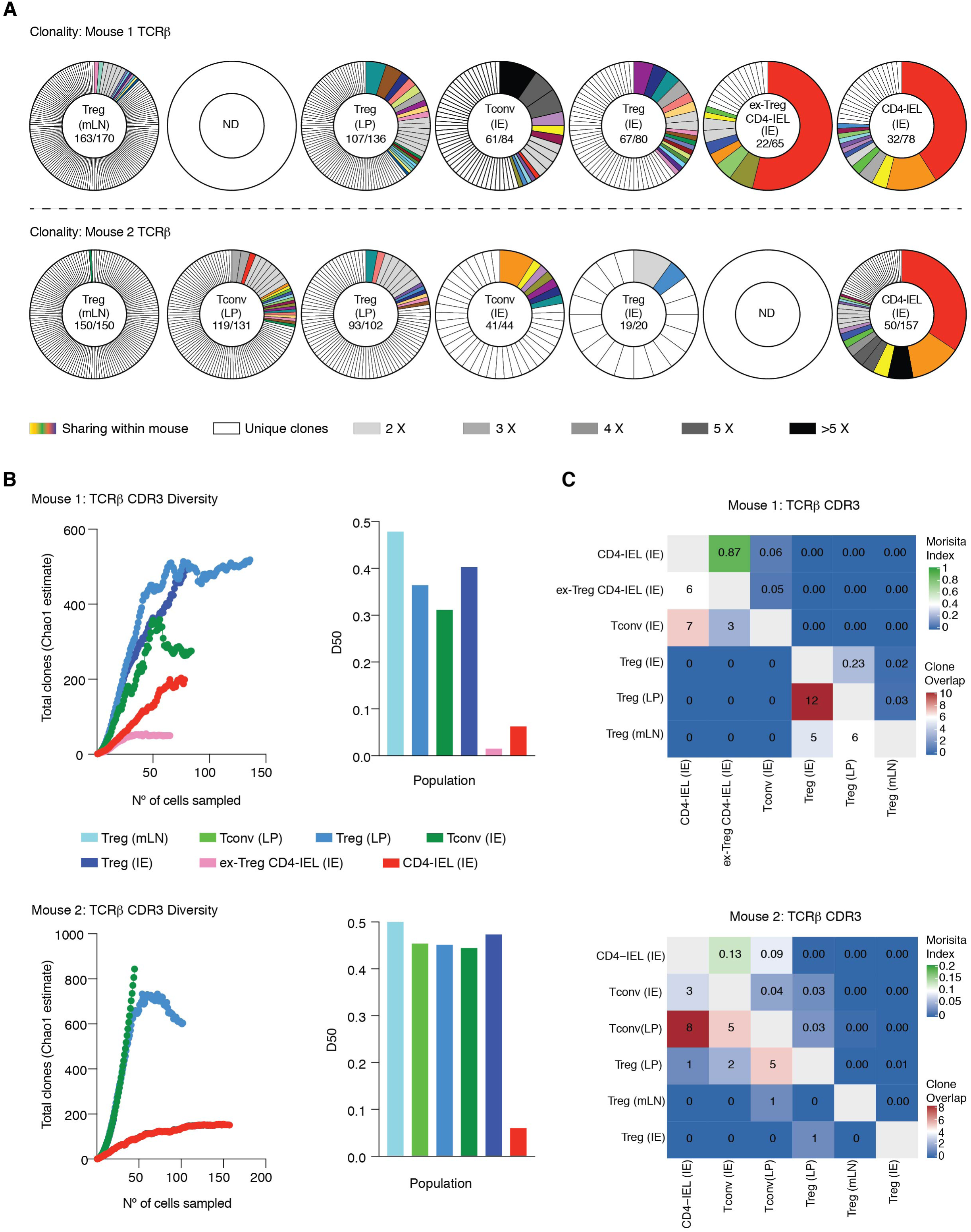
CD4-IELs are clonally expanded with decreased TCR diversity. (**A-C**) *Foxp3*^eGFP-Cre-ERT2^ x *Rosa26*^lsl-tdTomato^ x *Foxp3*^IRES-GFP^ (i*Foxp3*^Tom^ *Foxp3*^GFP^) mice were treated with tamoxifen for 10 weeks, and CD4^+^ T cells from mesenteric lymph nodes (mLN), lamina propria (LP) and intestinal epithelium (IE) were sorted as follows: CD4^+^ conventional (Tconv; GFP^−^Tomato^−^CD8α^−^), regulatory T cells (Treg; GFP^+^ or Tomato^+^CD8α^−^), ex-Treg CD4-IEL (Tomato^+^CD8α^+^) and CD4-IEL (GFP^−^Tomato^−^ CD8α^+^). TCRβ were sequenced via the MiSeq platform. TCRα of expanded TCRβ clones were also sequenced to confirm clonality. (**A**) TCRβ clonal diversity of indicated populations from two separate mice. Each slice represents a distinct TCRβ CDR3. Colored clones represent sharing within each mouse. White slices represent unique clones and grey-scale slices represent expanded clones at indicated populations. The numbers enclosed in each graph indicates number of clones (numerator) and total number of cells (denominator) per indicated population. Empty graphs with “ND” indicate no data for corresponding cell types. (**B**) Diversity estimated by Chao1 estimate (left) and D50 (right) of indicated cells in two separate mice based on their TCRβ CDR3s. (**C**) Normalized Morisita index (top right) and number of shared clones (bottom left) of TCRβ CDR3s per cell type of both mice.

We observed a diverse repertoire among Tregs isolated from mLN; from 320 sequenced cells, we retrieved 313 unique clones (Figure 1A, B). TCR repertoire of Tregs and Tconv isolated from LP and IE was also diverse (Figure 1A, B). In contrast, we observed large clonal expansions and reduced TCR diversity among CD4-IELs and ex-Treg CD4-IELs (Figure 1A, B). Additionally, we detected clonal sharing between LP Tregs or Tconv and all IEL subsets analyzed: Tconv, CD4-IEL or ex-Treg CD4-IEL (Figure 1A, C), suggesting that peripheral CD4^+^ T cells first migrate to the lamina propria before entering the epithelium or that the same T cell clones migrate simultaneously to both locations. Moreover, several expanded clones were shared between ex-Treg CD4-IELs (Tomato^+^) and CD4-IELs (Tomato^−^), including some with Tconv, raising the possibility that a single Tconv precursor can differentiate into Tregs and then to CD4-IELs, in addition to directly converting to CD4-IELs (Figure 1 A, C). Analysis of an additional i*Foxp3*^Tom^ mouse showed similar results (Figure S1C, D). Because of the low proliferation capacity of CD4-IELs (Mucida et al., 2013), these results indicate that potential precursors migrate to the epithelium and proliferate before they can fully differentiate into CD4-IELs.

### Clonal distribution follows the trajectory of CD4-IEL differentiation

Our previous live imaging and fate-mapping studies suggest that emigrating CD4^+^ T cells quickly acquire an IEL program at the gut epithelium while losing hallmarks of peripheral CD4^+^ T cells or Tregs, including the expression of ThPOK and Foxp3, respectively (Reis et al., 2013; Sujino et al., 2016). To concomitantly address intestinal epithelium-induced CD4^+^ T cell plasticity and specific TCR features in this process, we performed 5’ single-cell RNA sequencing (scRNAseq) coupled to TCRseq analysis using the Chromium Single Cell V(D)J platform (10X Genomics). This strategy also allowed analysis of intra-mouse TCR sharing between current Tregs and ex-Treg CD4-IELs, which was not possible in our dual Foxp3 reporter-fate-mapping strategy used above given that *Foxp3* is X-linked. We analyzed tamoxifen-treated i*Foxp3*^Tom^ mice by sorting all Tomato^+^ (library 1) or Tomato^−^ (library 2) CD4^+^CD8β^−^ T cells, therefore examining the whole spectrum of heterogeneity of CD4^+^ T cells that gained access to the epithelium. We confirmed the presence of CD4-IELs and Tregs in sorted cells by expression of CD8α and Foxp3, respectively (Figure S2A). From the two libraries, we obtained a total of 1,294 scRNAseq profiles (898 for Tomato^+^ library 1 and 396 for Tomato^−^ library 2) with paired αβTCR sequences for 952 cells (651 for Tomato^+^ library 1 and 301 for Tomato^−^ library 2).

We identified 8 clusters ordered by cell number (0-7) visualized by UMAP (Figure 2A, Figure S2B, Table S1), comparable to what we identified in our parallel unpublished study describing the molecular mechanisms of the differentiation towards CD4-IELs (London *et al*., unpublished). The cluster of cycling cells (7) was excluded from most of the downstream analysis due to low number of cells and strong proliferation gene signature that segregated them from all the other clusters. We identified three Treg clusters (3, 5 and 6) composed primarily of Tomato^+^ cells that express genes ascribed to Tregs (Figure S2C,F, Table S1). Of note, cluster 5 contains cells with a profile of resting Tregs of lymphoid origin (*Tcf7, Il7r* and *Ccr7)*, includes a small subset of *Sell*-expressing cells (Naïve*) and was classified as recent emigrant Treg (RE-Treg; Figure S2D, Table S1). In addition to *bona-fide* Treg genes (*Foxp3, Ikzf2 Capg*), cells within Treg cluster 3 also express a profile associated to non-lymphoid tissue Tregs (*Tnfrsf4, 9, 18* and *Tigit)* (Miragaia et al., 2019). A small fraction of cells in cluster 3 (TCR*) expressed high levels of *Nr4a* and *Egr* family members, related to increased TCR stimulation and activation (Zemmour et al., 2018). Treg-like (cluster 1) was ascribed to cells expressing a Treg profile, yet to a lower extent than the Tregs of cluster 3, while also expressing some IEL genes (*Cd7* and *Gzmb*) (Table S1, Figure S2E). Finally, we identified three non-Treg clusters, one of them composed mostly of Tomato^+^ cells (cluster 0), indicating Treg origin, while the other two (clusters 2 and 4) contained a mix of Tomato^+^ and Tomato^−^ cells (Figure 2A, Figure S2C, F, Table S1). Cluster 2 was rather homogeneous containing CD8α-expressing cells with a “full IEL program” (Cheroutre et al., 2011; McDonald et al., 2018), which included the expression of *Gzma, Gzmb, Cd244 (2B4)*,and *Itgae* (CD103) (Figure 2A, Figure S2C, Table S1). Trajectory analysis allowed the inference of a peripheral CD4^+^ T cell differentiation hierarchy as they gain access to the epithelium (Figure 2B). We observed three different potential trajectories, all leading to the differentiation of CD4-IELs from either Tregs (Tomato^+^) or from Tconv (Tomato^−^) (Figure 2B). Clusters 0 and 4 directly preceded the CD4-IEL cluster 2 and we refer to them as “pre-IEL1” (cluster 0) and “pre-IEL2” (cluster 4). Together, these findings suggest that CD4-IELs represent a final stage of differentiation, and that peripheral CD4^+^ T cells acquire a similar IEL program regardless of their origin (Tomato^+^ or Tomato^−^).

**Figure 2.**
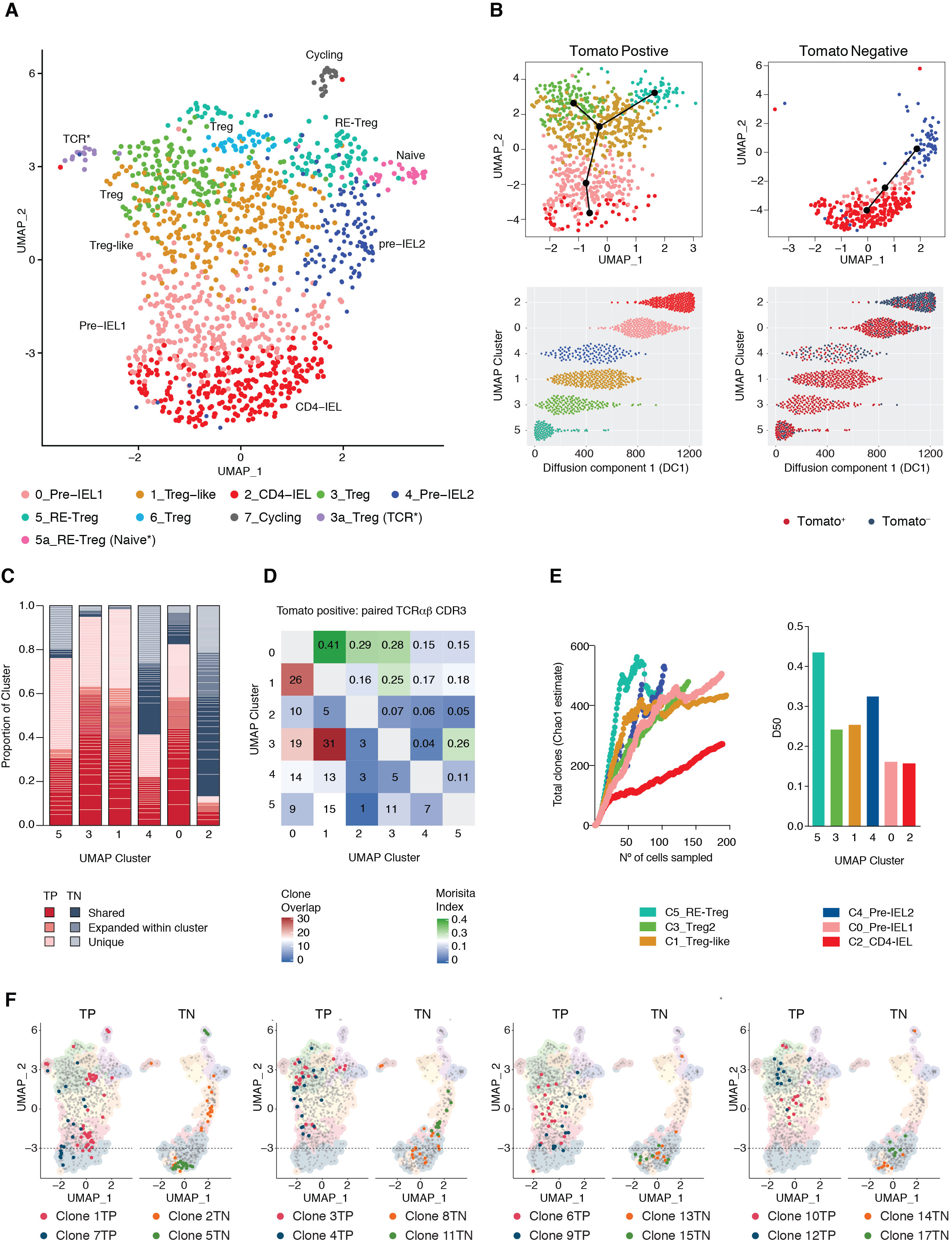
Clonal distribution of intraepithelial CD4^+^ T cells follows the single-cell trajectories. (**A-F**) i*Foxp3*^Tom^ mice were treated with tamoxifen for 10 weeks, and Tomato^+^ (library 1) and Tomato^−^ (library 2) CD4^+^ T cells from the intestinal epithelium (IE) were sorted for scRNAseq using 10X Genomics platform. (**A**) UMAP clustering of 8 (0-7) distinct populations of single cells, including sub-clusters (3a, 5a). Sub-cluster 3a_Treg (TCR*) indicates Tregs expressing TCR-stimulated genes. Sub-clusters 5a_RE-Treg (Naïve*) indicates *Sell*-expressing Tregs. Cluster names correspond to cell colors throughout the figure as indicated. (**B**) Pseudotime trajectory analysis of Tomato^+^ (top left) and Tomato^−^ (top right) cells. Cells on UMAP clusters ordered along diffusion component 1 (DC1), colored according to cluster (bottom left) or tomato expression (bottom right). (**C**) Paired αβTCR CDR3 of cells per UMAP cluster ordered by pseudotime trajectories of Tomato^+^ (TP, shades of red) and Tomato^−^ (TN, shades of blue) cells. Lightest shades indicate unique clones, intermediate shades indicate expanded, but not shared clones within TP or TN. Darkest shades of red or blue indicates αβTCR sharing between clusters per TP or TN, respectively. (**D**) Normalized Morisita index (top right) and absolute number of shared clones (Clone overlap, bottom left) of paired αβTCR per UMAP cluster among tomato positive cells. (**E**) Diversity estimated by Chao1 estimate (left) and D50 (right) of cells based on paired αβTCR per indicated cluster. (**F**) Top expanded clones per tomato positive (TP, red and blue) and negative (TN, orange and green) indicated in UMAP clusters, separated by tomato expression. Dashed line indicates top limit of CD4-IEL cluster 2.

Overlaying the TCRαβ repertoire information obtained from each cell on the UMAP clusters revealed variable levels of clonal expansion and spreading across the identified clusters (Figure 2C-F, Figure S2G,H). Consistent with our previous scTCRseq analysis, the CD4-IEL cluster was the least diverse cluster, as measured by the Chao1 index. Interestingly, although pre-IEL1 cluster 0 is not as diverse, it displayed similar level of clonal expansion, or dominance, as the CD4-IELs, as indicated by the low Diversity 50 (D50) score. Likewise, Treg and Treg-like clusters 3 and 1, respectively, displayed the next highest levels of clonal dominance (Figure 2E). Combination of pseudotime analysis (Figure 2B) with clonal distribution (Figure 2C, Figure S2G) and clonal diversity and dominance scores (Figure 2E), suggested that clonal expansion increased as cells developed towards CD4-IELs, but did not become homogeneous until the final CD4-IEL development stage. Furthermore, this analysis revealed a series of pathways through which different expanded clones differentiate from one state to another. This was especially true for fate-mapped cells, for which our tamoxifen labeling strategy served as a timestamp as Treg clones entered the epithelium, differentiated into pre-IELs and then into ex-Treg CD4-IELs. All of the top expanded clones that were present among the CD4-IELs were also found in at least one more cluster, with distributions that tended to follow the pseudotime trajectories (Figure 2C, D, F), suggesting expanded CD4^+^ T cells undergo differentiation at the epithelium. Our clonal composition analysis also indicates that terminal differentiation of CD4-IELs directly correlates with reduction in TCR diversity (Figure 2E). For example, the top expanded Treg-derived Tomato^+^ clones (clones 1TP and 7TP) were present in Treg, Treg-like, pre-IEL1 and CD4-IEL clusters, while some other expanded clones (clones 3TP, 4TP, 6TP, 9TP,10TP, 12TP) were not found among CD4-IELs but were shared between Tregs, Treg-like and pre-IEL1 clusters. Of note, the top expanded Tconv-derived Tomato^−^ clones were found within pre-IEL2 and CD4-IEL clusters (Figure 2F, Table S2). It is possible that a fraction of T cell precursors did not receive sufficient signals to convert into CD4-IELs, or that the time required for conversion was longer than the timeframe of our analysis. Overall, our findings reveal a high degree of TCR sharing between rather heterogeneous gut CD4^+^ T cell populations, with reduced diversity as cells differentiate into IELs.

### Decreased TCR signaling precedes IEL differentiation

Analysis of the scRNAseq data showed that acquisition of IEL markers, such as *Itgae* (CD103) by CD4^+^ T cells in the epithelium, is inversely correlated with expression of genes downstream of TCR signaling and co-stimulation, such as *Nr4a1* (Nur77) and *Tnfrsf4* (OX40). As pre-IELs acquire IEL and cytotoxic markers, they downmodulate TCR signaling molecules (Figure 3A, B). To confirm that downstream TCR signaling is associated with peripheral IEL differentiation, we analyzed Nur77 expression along differentiating peripheral IELs using *Nur77*^GFP^ *Foxp3*^RFP^ reporter mice (Figure S3 A-C). CD4-IELs and CD4^+^CD103^+^ cells, which are enriched in pre-IELs, express lower levels of Nur77 when compared to recently-emigrated CD4^+^CD103^−^ cells or to Tregs found in the epithelium (Figure 3C). Thus, Nur77 expression is inversely associated with acquisition of CD103 and CD8αα by CD4^+^ T cells in the epithelium, suggestive of a role for TCR signaling in IEL differentiation from peripheral CD4^+^ T cells.

**Figure 3.**
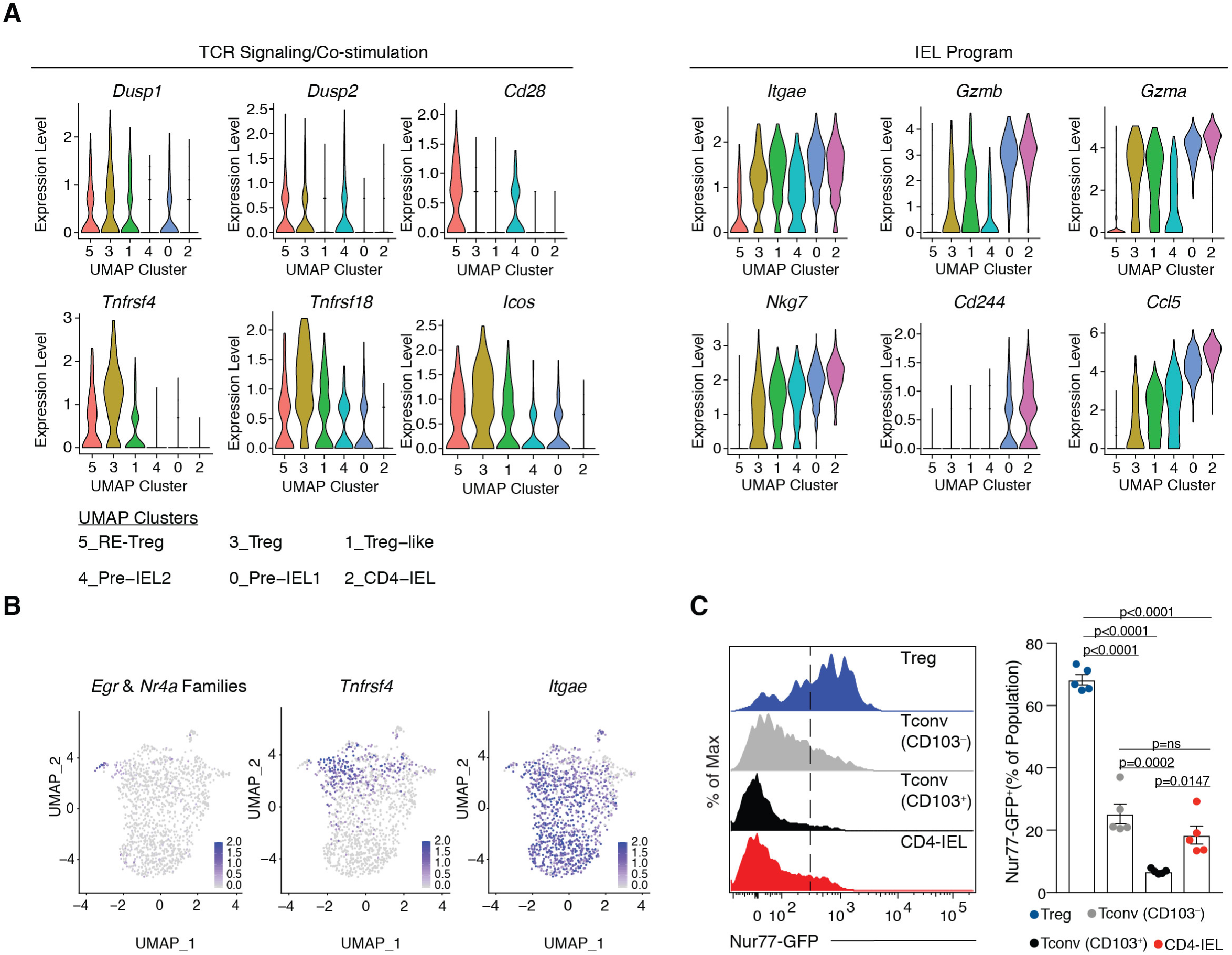
Intraepithelial CD4^+^ T cell subsets show an inverse correlation between TCR signaling and IEL program. (**A, B**) i*Foxp3*^Tom^ mice were treated with tamoxifen for 10 weeks, and Tomato^+^ (library 1) and Tomato^−^ (library 2) CD4^+^ T cells from the intestinal epithelium (IE) were sorted for scRNAseq using 10X Genomics platform. (**A**) Expression levels of genes related to TCR signaling or IEL program in each UMAP cluster ordered by pseudotime trajectories. (**B**) Expression levels of *Egr* and *Nr4a* families (left), *Tnfrsf4* (OX40, middle) and *Itgae* (CD103, right) in all sequenced cells. (**C**) Nur77 as measured by GFP fluorescence expression levels (left) and frequencies (right) among Foxp3^+^ regulatory T cells (RFP^+^, Treg, blue), conventional CD4^+^ T cells (RFP^−^, CD8α^−^, Tconv) CD103^−^ (grey) or CD103^+^ (black) cells, and CD4-IELs (RFP^−^, CD8α^+^TL-Tetramer^+^, red) in the small intestinal epithelium of SPF *Nur77*^GFP^*Foxp3*^RFP^ double-reporter mice.

### TCR signaling is required for CD4-IEL development

To directly assess the requirement of the TCR for CD4-IEL differentiation, we employed multiple Cre–mediated TCR ablation strategies targeting different stages or subsets of CD4^+^ T cells. First, we addressed whether TCR signaling is required for Treg plasticity in the gut epithelium and subsequent CD4-IEL differentiation by crossing *Trac*^f/f^ mice to i*Foxp3*^Tom^ mice (to generate the i*Foxp3*^TomΔ(*Trac*)^ strain), which allowed for the tracking of ex-Tregs as they lose surface TCR expression (Figure 4A,B, Figure S4A,B). Whereas CD4-IEL differentiation from TCR-expressing (TCRβ S^+^; Tomato^−^ or Tomato^+^) cells was similar between i*Foxp3*^Tom^ and i*Foxp3*^TomΔ(*Trac*)^ mice, it was significantly impaired among TCR-deficient (TCRβ S^−^ Tomato^+^) cells of i*Foxp3*^TomΔ(*Trac*)^ mice (Figure 4A, B, Figure S4C). Our scRNAseq data showed that *Tnfrsf4* (OX40) expression changed in a developmentally controlled manner, peaking in Treg cluster 3 and then rapidly decaying by pre-IEL clusters 4 and 0 (Figure 3A, B). This pattern of expression allowed us to use the well-established OX40^Cre^ driver (Reis et al., 2013) to delete the TCR from IEL precursors but not from IELs themselves. We confirmed TCR deletion in activated CD4^+^ T cells and Tregs in the mLN of *Trac*^f/f^ x OX40^Cre^ (OX40^Δ(*Trac)*^) mice by flow cytometry (Figure S4F-J). In the epithelium, while the total frequency of CD4^+^ T cells remained the same in OX40^WT(*Trac)*^ and OX40^Δ(*Trac)*^ mice (Figure S4D), TCR ablation significantly decreased the CD4-IEL population (Figure 4C, D, Figure S4E). We further compared the frequencies of CD4-IELs among TCR-sufficient and TCR-deficient CD4^+^ T cells in OX40^Δ(*Trac)*^ mice based on surface TCRβ expression. CD4-IELs were only observed among TCR-sufficient cells (Figure 4E). Taken together, our data indicate that the differentiation of both Treg and Tconv to CD4-IELs requires TCR expression.

**Figure 4.**
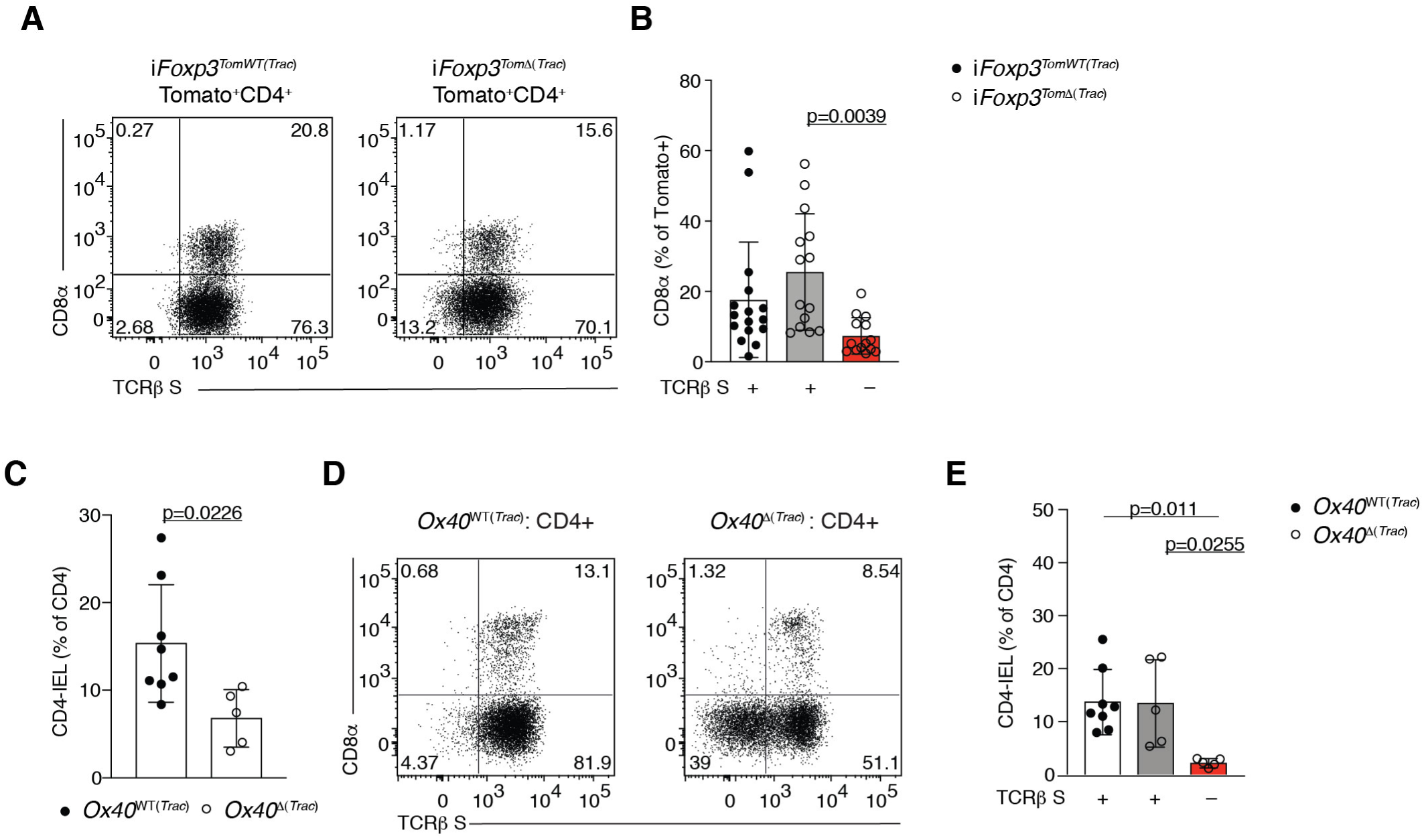
TCR signaling is required for CD4-IEL differentiation. (**A, B**) Flow cytometry analysis of the intestinal epithelium (IE) of i*Foxp3*^*Tom*WT*(Trac)*^ (*Trac*^+/+^ i*Foxp3*^Tom^) or i*Foxp3*^*Tom*Δ(*Trac)*^ (*Trac*^f/f^ i*Foxp3*^Tom^) mice 8-12 weeks after tamoxifen administration. (**A**) Representative dot plots of surface CD8α and TCRβ of Tomato^+^ CD4^+^ T cells in i*Foxp3*^*Tom*WT*(Trac)*^ (left) and i*Foxp3*^*Tom*Δ(*Trac)*^ (right) animals. (**B**) Frequencies of CD8α^+^ cells among Tomato^+^ CD4^+^ T cells within TCR-sufficient cells from i*Foxp3*^*Tom*WT*(Trac)*^ (white bar) or i*Foxp3*^*Tom*Δ(*Trac)*^ (grey bar) mice, or TCR-deficient cells from i*Foxp3*^*Tom*Δ(*Trac)*^ (red bar) mice. (**C-E**) Flow cytometry analysis of CD4^+^ T cells in the IE of 9-12-week-old OX40^WT(*Trac)*^ (*Trac*^+/+^ OX40Cre^+/−^ or *Trac*^f/f^ OX40Cre^−/−^) or OX40^Δ(*Trac)*^ (*Trac*^f/f^ OX40Cre^+/−^) mice. (**C**) Frequency of CD4-IELs (CD8α^+^TL-Tetramer^+^) among CD4^+^ T cells. (**D**) Representative dot plots of surface CD8α and TCRβ of CD4^+^ T cells. (**E**) Frequencies of CD4-IELs among conventional CD4^+^ T cells (Tconv, CD4^+^CD8α^−^Foxp3^−^) within TCR-sufficient cells from OX40^WT(*Trac)*^ (white bar) or OX40^Δ(*Trac)*^ (grey bar) mice, or TCR-deficient cells from OX40^Δ(*Trac)*^ (red bar) mice. Data are expressed as mean +/− SEM of individual mice (n=5-16). Significant p values as indicated [student’s t test (**C**) or one-way ANOVA and Bonferroni (**B, E**)].

### MHCII expression on epithelial cells modulates CD4-IEL differentiation

Recent studies have suggested a role for intestinal epithelial cell (IEC)-mediated antigen presentation via MHCII in the regulation of intestinal CD4^+^ T cell function (Biton et al., 2018; Koyama et al., 2019). We therefore asked whether local MHCII expression by IECs is required for CD4-IEL differentiation or maintenance. We targeted MHC class II expression exclusively on IECs by crossing Villin^CreERT2^ mice to *H2-Ab1*^f/f^ mice (Villin^Δ(MHCII)^) (Figure S5A-C). Tamoxifen treatment of Villin^Δ(MHCII)^ mice starting at 5-7 weeks of age (prior to the appearance of CD4-IELs in the epithelium) did not affect the frequency of total CD4^+^ T cells in the epithelium 5-6 weeks later; however, it led to a significant reduction in the frequency of CD4-IELs, which was accompanied by an increase in Treg frequency (Figure 5A). Tamoxifen treatment of 11, 12- or 16-week-old mice, which typically carry a sizable population of CD4-IELs, also significantly impacted CD4-IEL frequencies and conversely led to an accumulation of Tregs in the epithelium (Figure 5B, C), suggesting that MHCII expression on IECs is required for continuous differentiation into CD4-IELs in adult mice.

**Figure 5.**
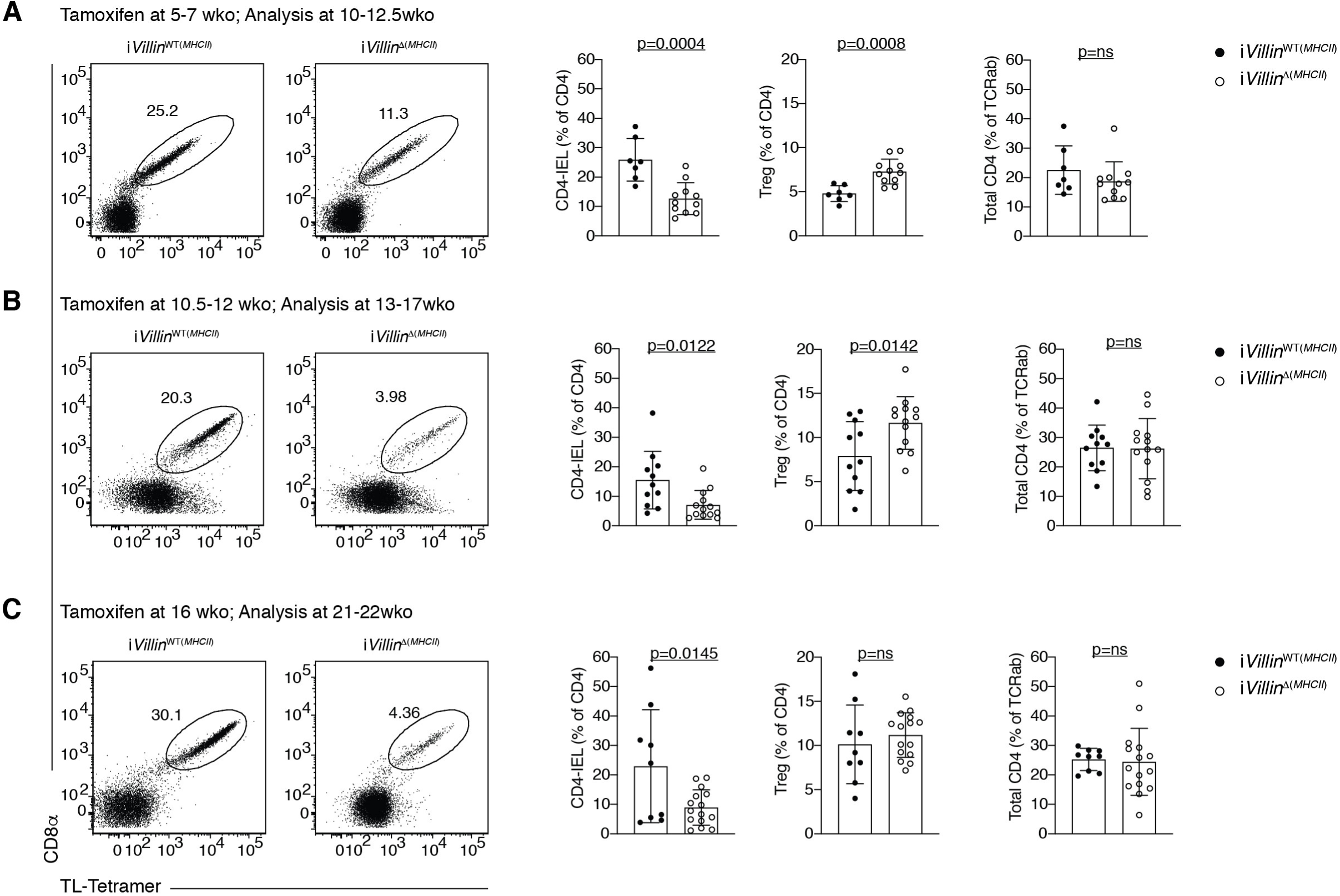
MHCII expression by epithelial cells is required for CD4-IEL conversion. (**A-C**) Flow cytometry analysis of the intestinal epithelium (IE) of i*Villin*^WT(*MHCII)*^ (*H2-Ab1*^+/+^ Villin^CreERT2+/–^ or *H2-Ab1*^f/f^ Villin^CreERT2–/–^) or i*Villin*^Δ(*MHCII)*^ (*H2-Ab1*^f/f^ Villin^CreERT2+/+^) mice after tamoxifen administration to 5-7 week-old mice, analyzed at 10-12.5 weeks (**A**), to 10.5-12 week-old mice, analyzed at 13-17 weeks of age (**B**), to 16 week-old mice, analyzed at 21-22 weeks of age (**C**). Representative dot plots of surface CD8α and TL-Tetramer among CD4^+^ T cells (left). Frequencies of CD4-IELs (CD4^+^CD8α^+^TL-Tetramer^+^) or Foxp3^+^ regulatory cells (Tregs) among CD4^+^ T cells (middle), and total CD4^+^ T cells among TCRαβ^+^ cells (right). Data are expressed as mean +/− SEM of individual mice (n=9-15). Significant p values as indicated [student’s t test (**A-C)**].

### TCR signaling is largely dispensable for IEL program maintenance

The transition to an IEL program includes upregulation of NK- and cytolytic-molecules as well as CD8αα (Cheroutre et al., 2011; McDonald et al., 2018), with concomitant downregulation of TCR signaling. To evaluate the role of the TCR signaling in establishing the IEL program, and in the maintenance of differentiated IELs, we crossed *Trac*^f/f^ mice to those expressing Cre under the enhancer I of the *Cd8a* gene (E8_I_^Δ(*Trac)*^). E8_I_ is required for CD8α expression on mature T cells and CD8αα IELs, but not required for CD8αβ or CD8αα expression in developing thymocytes (Ellmeier et al., 1997). In E8_I_^Δ(*Trac)*^ mice, TCR deletion is accompanied by a decrease in CD8α-expressing TCRαβ^+^ cells in the mLN and IE, including CD4^−^CD8β^+^CD8α^+^ T cells and CD4^−^CD8β^−^CD8αα^+^ TCRαβ^+^ natural IELs (nIEL) (Figure S6A-F). In the case of peripheral CD4^+^ T cells, which only express CD8αα in the final stages of their differentiation into CD4-IEL, this model allowed us to selectively address the role of the TCR in the maintenance of differentiated CD4-IELs. We did not observe any significant changes in the accumulation of CD4-IELs in E8_I_^Δ(*Trac)*^ mice when compared to WT littermates (E8_I_^WT(*Trac)*^), even in older animals (Figure 6A,B), suggesting that upon terminal differentiation, CD4-IELs do not rely on the TCR for their maintenance in the epithelium. Whereas proliferation, measured by Edu incorporation, was similar among Tconv from E8_I_^WT(*Trac)*^ and E8_I_^Δ(*Trac)*^ mice, remaining TCR-expressing CD4-IELs in E8_I_^Δ(*Trac)*^ mice proliferated more than those of E8_I_^WT(*Trac*)^ mice, but only in older animals (Figure 6C, D). Therefore, similar frequencies of CD4-IELs cannot be exclusively attributed to differential proliferative abilities of TCR-sufficient and TCR-deficient cells. Similar proliferation rates were observed in other CD8αα-expressing IEL subsets, including CD8αβ^+^CD8αα^+^ TCRαβ^+^ (CD8-IELs) and CD8αα^+^ nIELs (Figure S6G, H). Common IEL functional readouts such as IFNγ production and granzyme B expression showed that hallmarks of the CD4-IEL phenotype are maintained in the absence of the TCR (Figure 6E, F). Analysis of CD8-IELs and nIELs also revealed intact IFNγ and granzyme B production, despite TCR loss (Figure S6I-L). Taken together, our results suggest that CD4-IEL accumulation and at least some of the characteristic features of IELs are maintained in the absence of TCR signaling.

**Figure 6.**
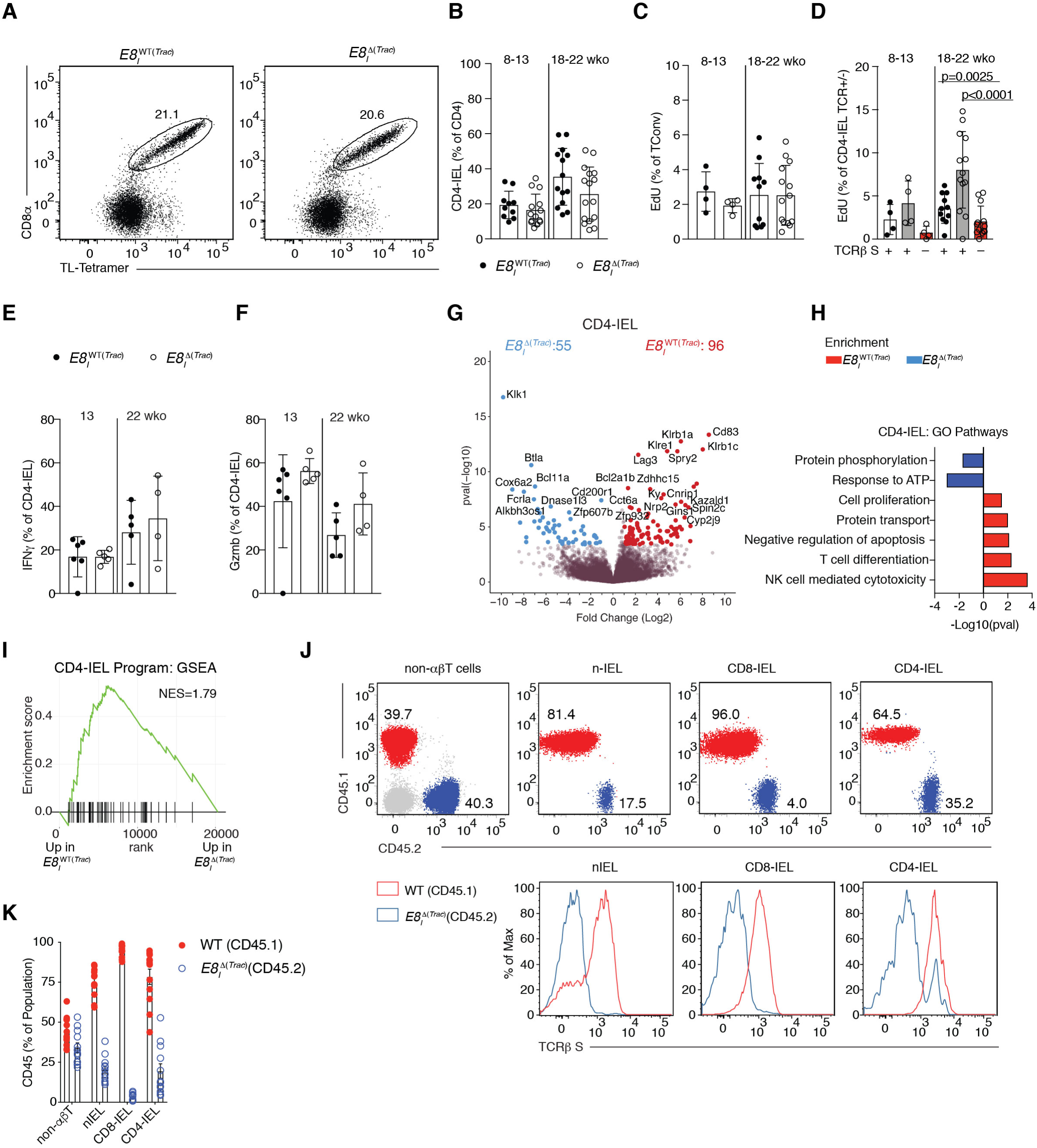
TCR signaling is not essential for CD4-IEL maintenance. (**A-F)** Flow cytometry analysis of intestinal epithelium (IE) of 8-22 week-old E8_I_^WT(*Trac)*^ (*Trac*^+/+^ E8_I_Cre^+^ or *Trac*^+/+^ E8_I_Cre^−^ or *Trac*^f/f^ E8_I_Cre^−^) or E8_I_^Δ(*Trac)*^ (*Trac*^f/f^ E8_I_Cre^+^) mice, grouped by age as indicated. (**A**) Representative dot plots of surface CD8α and TL-Tetramer among CD4^+^ T cells in E8_I_^WT(*Trac)*^ (left) or E8_I_^Δ(*Trac)*^ (right) mice. (**B**) Frequency of CD4-IELs (CD8α^+^TL-Tetramer^+^) among CD4^+^ T cells in E8_I_^WT(*Trac)*^ and E8 _I_^Δ(*Trac)*^ mice. (**C, D**) Frequency of proliferation (measured by EdU incorporation) of CD4^+^CD8α^−^ Foxp3^−^ cells (Tconv) (**C**) among cells with or without surface TCR expression (**D**). Mice were injected with Edu 16 and 4 hours prior to analysis. (**E, F**) Frequencies of IFNγ (**E**) and Gzmb (**F**) production upon PMA/Ionomycin *ex-vivo* stimulation among CD4-IELs. (**G-I**) Bulk RNA-sequencing was performed on TCRβ^+^ CD4-IELs from E8_I_^WT(*Trac)*^ and TCRβ^−^ CD4-IELs from E8_I_^Δ(*Trac)*^ mice (n=3 per group). Volcano plot of differentially expressed genes between indicated populations (p<0.05, in color) (**G**), selected differentially-enriched gene ontology (GO) pathways between groups (**H**), and gene set enrichment analysis (GSEA) of CD4-IEL program as defined by the top differentially-expressed genes in the CD4-IEL cluster in our single cell RNA-Sequencing from Figure 2 (**I**). (**J, K**) Flow cytometry analysis of bone marrow chimeras in sub-lethally irradiated *Rag1*^*–/–*^ hosts reconstituted with 1/1 ratio of WT CD45.1 and E8_I_^Δ(*Trac)*^ CD45.2 cells, analyzed 12-16 weeks after reconstitution. Representative dot plots and histograms (**J**) and frequency (**K**) of WT CD45.1 (red) versus E8_I_^Δ(*Trac)*^ CD45.2 (blue) among non-αβT cells (includes γδT cells and non-T cells), natural IELs (nIEL, CD4^−^CD8αα^+^CD8β^−^TL-Tetramer^+^), CD8-IELs (CD4^−^CD8αα^+^CD8β^+^TL-Tetramer^+^), and CD4-IELs in the IE. Bottom panels, representative histograms of surface TCRβ levels in WT CD45.1 (red line) and E8 _I_^Δ(*Trac)*^ CD45.2 (blue line) of indicated cell populations. Frequency data are expressed as mean +/− SEM (**B-F**). Significant p values as indicated [student’s t test (**B, C, E, F**) or one-way ANOVA and Bonferroni (**D**)]. n=10-16 (**B**), n=4-14 (**C, D**), n=4-6 (**E, F**) and n=12 (**K**).

To comprehensively address the extent to which the IEL program can be affected or maintained in the absence of TCR signaling, we performed bulk RNAseq on TCR-expressing CD4-IELs from E8_I_^WT(*Trac)*^ mice and on TCR-deficient CD4-IELs from E8_I_ ^Δ(*Trac)*^ mice (Figure S6M). While genes related to TCR activation or signaling, cell cycle or proliferation, anti-apoptosis and protein transport showed reduced expression in cells lacking their TCR, most IEL-related genes were not significantly changed in E8_I_^Δ(*Trac)*^ mice, suggesting the IEL program is maintained in the absence of TCR expression (Figure 6G-I, Table S3). Gene ontology (GO) enrichment analysis showed that only a few pathways, such as protein phosphorylation, were upregulated in TCR-deficient cells (Figure 6H). Likewise, IEL genes were not changed among CD8-IELs or nIELs in E8_I_^Δ(*Trac)*^ mice, but genes related to TCR signaling were significantly decreased in TCR-deficient cells (Figure S6 N,O, Table S4,5). Of note, Lag3 (Lymphocyte activation gene-3), a TCR inhibitory co-receptor, was downmodulated in all 3 IEL subsets lacking TCR expression (Figure 6G, Table S1).

To further evaluate whether the TCR confers any competitive advantage to IELs, we reconstituted sub-lethally irradiated *Rag1*^−/−^ mice with a 1:1 mix of congenically-marked bone marrow from WT CD45.1 and E8_I_^Δ(*Trac)*^ CD45.2 mice (Figure 6J). Whereas the ratio of WT to E8_I_^Δ(*Trac)*^ remained approximately 1:1 in non-αβT cells, this ratio was significantly skewed towards WT among CD8-IELs (Figure 6J,K), indicating that TCR-deficient CD8-IELs were outcompeted by TCR-expressing cells. In contrast, within both nIELs and CD4-IELs, the WT: E8_I_^Δ(*Trac)*^ ratio was much less skewed, and surface-TCRβ^−^ cells were readily detected among these populations (Figure 6J). Thus CD4-IELs are much less sensitive than CD8-IELs to TCR loss. Together, our data establish an important role of TCR in IEL differentiation, particularly in the final stages within the epithelium, but also indicate that TCR signaling is mostly dispensable for CD4-IEL maintenance.

## Discussion

Self-reactive T cell populations with regulatory properties such as natural Tregs, iNKT cells, and CD8αα IELs are selected via agonist-selection, a process that favors development of cells with strong TCR avidity despite of self-reactivity (Sakaguchi et al., 2013; Yamagata et al., 2004). However, much less is known about how TCR signaling and repertoire impact tissue-induced plasticity and the differentiation of peripheral T cells which are presumably selected on weaker but broader TCR affinity interactions.

Our study revealed a previously unappreciated degree of TCR sharing between largely distinct CD4^+^ T cell subsets in the periphery, distinct levels of clonal expansion and TCR diversity along differentiating IELs, and a specific requirement for TCR signaling during the early stages of the IEL differentiation process.

Early analyses of TCR diversity in nIEL subsets revealed a restriction in TCR repertoire, referred to as “oligoclonal repertoire” (Guy-Grand et al., 1991; Regnault et al., 1994; Regnault et al., 1996; Rocha et al., 1991). More recently, work using a TCR^mini^ mouse model harboring restricted TCRα and TCRβ repertoires showed a substantial TCR overlap between CD4-IELs and Tregs (Wojciech et al., 2018). In addition to the agonist selection of thymic IEL precursors (Leishman et al., 2002; Yamagata et al., 2004), studies that generated transgenic mouse strains carrying existing IEL αβTCRs strongly suggested that TCR specificity may be sufficient to drive IEL fate (Mayans et al., 2014; McDonald et al., 2014). It remained unclear, however, how specific TCRs correlate with IEL differentiation. Our scRNAseq and trajectory analyses coupled to TCR repertoire allowed us to unbiasedly define the relationship between TCR diversity and CD4^+^ T cell plasticity during migration and differentiation towards IELs. Expanded clones followed pseudotime trajectory analysis and displayed intra-clonal plasticity: less expanded clones spread among heterogenous subsets, while highly expanded clones were found in the homogenous cluster of CD4-IELs. A possible explanation for this finding is that the less expanded clones lacked additional signals, such as TCR ligands or environmental components, required for full differentiation into CD4-IELs (Cervantes-Barragan et al., 2017; Cortez et al., 2014; Mucida et al., 2013; Reis et al., 2014; Reis et al., 2013). The clonal analyses presented here, including the TCR sharing of CD4-IELs derived from Treg or from Tconv, corroborate previous studies suggesting a lineage relationship between Tregs and IELs (Bilate et al., 2016; Sujino et al., 2016). Additionally, their clonal distribution suggest that IEL differentiation may favor particular TCR specificities, perhaps in a process analogous to peripheral Treg differentiation, where TCR recognition in a context–dependent manner leads to Foxp3 expression (Curotto de Lafaille et al., 2004; Mucida et al., 2005).

Despite of being “chronically activated” as defined by the expression of activation markers such as CD69 and CD44 (Mucida et al., 2013), CD4-IELs do not show signs of strong TCR activation. We find that Nur77, which is expressed proportionally to the strength of the TCR stimulation, is increased in migrating CD4^+^ T cells but is progressively down-modulated as they differentiate into pre-IELs. This is similar to the high level of Nur77 displayed by agonist-selected T cells such as iNKT cells during thymic selection, which subsequently decays upon migration of these cells to the spleen or liver (Moran et al., 2011). While the majority of CD4-IELs express low levels of Nur77, a small fraction of them maintained Nur77 expression, suggesting that TCR re-engagement can be modulated within the epithelium. Indeed, availability of antigens presented by IEC could function in the late-stage of IEL differentiation, possibility supported by the decreased CD4-IEL population upon MHCII targeting on IECs. The modulation of MHCII expression by IECs has been linked to the capacity of microbes to attach to the epithelium, and to IFNγ production in both human and murine models (Ivanov et al., 2009; Panja et al., 1998; Umesaki et al., 1995). MHCII expression by IECs has been recently associated with a variety of physiological, such as regulation of the stem cell niche, and pathological functions, such as CD4^+^ T cell-mediated inflammation during graft-versus-host disease, in part through interacting with gut resident T cells that provide cytokines (Biton et al., 2018; Koyama et al., 2019; Ladinsky et al., 2019). Our data suggests a model in which antigen presentation by IECs is instrumental for the differentiation of CD4-IELs, which may further enhance MHCII expression by IECs via IFNγ production. Whether this is exclusively dependent on antigen presentation by IECs needs further demonstration. Regardless, the TCR complex itself, and presumably antigenic stimulation, is important for CD4-IEL differentiation as demonstrated by TCR ablation on OX40-expressing cells as well as on Tregs, which then precluded the differentiation into CD4-IELs.

However, our results show that TCR ablation on CD4-IELs does not impair their persistence in the epithelium nor the production of IFNγ and granzyme B. This is consistent with the decreased antigen sensitivity and increased threshold for TCR activation in cells expressing CD8αα homodimers (Cheroutre and Lambolez, 2008). Furthermore, our RNAseq analysis suggests that despite the down-modulation of genes downstream of TCR activation in TCR-deficient CD4-IELs, the maintenance of an IEL program may not depend on continuous TCR signaling. This is in contrast to the requirement of TCR expression on mature Tregs for their suppressive function and maintenance of an effector Treg program (Levine et al., 2014). The extent to which CD4-IELs depend on TCR signaling for specific functions and whether it is needed in discrete modules during different stages of differentiation from Tregs to CD4-IELs remains to determined.

Although surface expression of T cell receptors did not significantly impact CD4-IEL maintenance, TCR signaling and local MHC class II expression on IECs are essential for T cell plasticity at the epithelium. Given the potential abundance of antigens to which the intestinal epithelium is exposed, the restricted TCR diversity of CD4-IELs is intriguing. It is likely that CD4-IELs recognize a restrict set of peptide-MHCII complexes. The origins (microbial or dietary), and the range of TCR specificities and affinities for these antigens recognized by CD4-IELs still remains to be elucidated. This will help address the impact of MHC class II-restricted immune responses at the intestinal epithelium.

## Acknowledgements

We are grateful to A. Rogoz, J. Bortolatto and S. Gonzalez for exceptional animal care, mouse colony management and genotyping and the Rockefeller University employees for continuous assistance. We thank K. Gordon and K. Chhosphel for assistance with cell sorting. We thank C. Zhao and the entire Genomics Core of Rockefeller University for library preparation for 10X Genomics and assistance with all sequencing platforms used in this paper. We thank the NIH tetramer core facility for providing the TL tetramers used in this study. We are grateful to J. Lafaille for support throughout this project. We thank B. Reis for suggestions and critical reading of the manuscript, and all the members of the Mucida lab for fruitful discussions.

## Author contribution

AMB, ML and DM conceived the study, designed experiments and wrote the manuscript. AMB and ML performed and analyzed experiments. TBRC performed all bioinformatics analyses and assisted with interpretation of sequencing data. AH and SK helped with single-cell PCRs and multiplexing of samples for scTCRseq. LM helped with analysis of scTCRseq by Miseq, library preparation of bulk RNAseq and with calculation of diversity index, under the supervision of GDV.

## Methods

### Animals

Animal care and experimentation were consistent with NIH guidelines and were approved by the Institutional Animal Care and Use Committee at the Rockefeller University. *Rag1*^−/−^ (002216), *Thpok*^eGFP^ (027663), *Rosa26*^lsl-tdTomato^ (007914), *Foxp3*^eGFP-CreERT2^ (016961), CD45.1(B6.SJL *Ptprc*^a^, 002014), *Nr4a1*^EGFP/Cre^ (Nur77^GFP^, 016617), *Foxp3*^IRES-mRFP^ (008374), OX40^IRES-Cre^ (012839) mice were purchased from Jackson Laboratories and housed in our facility. *Trac*^f/f^ mice were kindly provided by A. Rudensky (MSKCC). *Villin*^CreERT2^ mice were generated by (el Marjou et al., 2004) and kindly provided by D. Artis (Cornell). *Foxp3*^IRES-GFP^ mice were provided by V. Kuchroo (Harvard) and *H2-Ab1*^f/f^ mice were provided by M. Nussenzweig (Jax 013181). E8_I_^Cre^ were kindly provided by I. Taniuchi (Jax 008766). Several of these lines were interbred in our facilities to obtain the final strains described elsewhere in the text. Genotyping was performed according to the protocols established for the respective strains by Jackson Laboratories or by donor investigators. Mice were maintained at the Rockefeller University animal facility under specific pathogen-free (SPF) conditions.

### Antibodies and additional flow cytometry reagents

Fluorescent dye–conjugated antibodies were purchased from BD Biosciences, Biolegend or Ebioscience (Thermofisher). The following clones were used: anti-CD45.1, A20; anti-Foxp3, FJK-16s; anti-CD4, RM4-5; anti-CD19, eBio1D3; anti-IFN-γ, XMG1.2; anti-CD45.2, 104; anti-CD8α, 53-6.7; anti-CD8β, YTS 156.7.7; anti-CD44, IM7; anti–I-A/I-E, M5/114.15.2; anti-CD45, 30-F11; anti-Granzyme B, NGZB; anti-CD103, 2E7; anti-CD62L, MEL-14; anti-EpCAM, G8.8; anti-TCRβ, H57-597; anti-TCRγδ, eBioG23. Live/dead fixable dye Aqua and Edu (ThermoFisher Scientific) were used according to manufacturer’s instructions. TL-Tetramer was obtained from NIH tetramer facility.

### Isolation of intestinal T cells

Intraepithelial and lamina propria lymphocytes were isolated as previously described (Bilate et al., 2016; Reis et al., 2013). Briefly, small intestines were harvested and washed in PBS and 1mM dithiothreitol (DTT) followed by 30 mM EDTA. Intraepithelial cells were recovered from the supernatant of DTT and EDTA washes and mononuclear cells were isolated by gradient centrifugation using Percoll. Lymphocytes from lamina propria were obtained after collagenase digestion of the tissue. Single-cell suspensions were then stained with fluorescently-labeled antibodies for 25min at 4°C prior to downstream flow cytometry (analysis or sorting) as specified in figure legends.

### Flow cytometry and intranuclear/intracellular staining

Flow cytometry data was acquired on an LSR-II flow cytometer (Becton Dickinson, USA) and analyzed using FlowJo software package (Tri-Star, USA). Intranuclear staining of Foxp3 was conducted using Foxp3 Mouse Regulatory T Cell Staining Kit (eBioscience, USA). For analysis of cytokine-secretion, IELs were plated in 48-well plates and incubated at 37°C with 100ng/mL phorbol 12-myristate 13-acetate (PMA, Sigma) and 200ng/mL ionomycin (Sigma) for 4 hours. Monesin (2μM, Sigma) was added 1h after PMA and ionomycin. Intracellular staining for IFNγ and granzyme B was conducted in Perm/Wash buffer after permeabilization in Fix/Perm buffer (BD Pharmingen, USA) according to kit instructions.

### Staining Strategy

For analysis of all Δ*Trac* strains (and their controls), the following gating strategy was utilized to examine CD4^+^ T cells: single live lymphocytes (based on size and live/dead stain), CD45^+^, TCRγδ^−^, intracellular TCRβ^+^, CD8β^–/low^, CD4^+^. For single-cell sorting of cells subjected to scTCRseq the following gating strategy was used: single live lymphocytes, CD45^+^, TCRγδ^−^, TCRβ^+^, CD8β^–/low^, CD4^+^, Tomato^+/−^ CD8α^+/−^ For sorting of cells subjected to bulk RNAseq we used single live lymphocytes CD45^+^, TCRγδ^−^, TCRβ^+/−^, CD8β^+/−^, CD4^+/−^, TL-Tetramer^+/−^ CD8α^+/−^ as indicated elsewhere in the text. For sorting of cells subjected to 10X Genomics, we gated on single live lymphocytes CD45^+^ TCRγδ^−^, TCRβ^+^, CD8β^–/low^, CD4^+^, Tomato^+/−^. IEL and Treg populations were confirmed by post-sort staining for surface CD8α and intranuclear Foxp3, respectively.

### Edu treatment and Detection

1mg EdU was injected intravenously at 5mg/mL in PBS 16 and 4 hours prior to analysis. Detection was performed using the Click-iT™ Plus EdU Flow Cytometery Assay kit (Thermo Fisher Scientific, C10632), according to manufacturer’s instructions.

### Tamoxifen treatment

Tamoxifen (Sigma) was dissolved in corn oil (Sigma) and 10% ethanol and shaking at 37°C for 30min-1h. Five doses of Tamoxifen (1mg/dose) was administered to *Villin*^CreERT2^ animals intraperitonially at 10mg/mL during one week, with 2 extra 1mg boosts 3 days apart 2 weeks before analysis, when analysis was more than 4 weeks after initial dosing, as indicated in figure legends. Four doses of Tamoxifen (5mg/dose) were administered to *Foxp3*^eGFP-CreERT2^ animals intragastrically 3 times 2 days apart in the first week, and then 2 times (3 days apart) every other week up to 2 weeks before analysis as indicated in figure legends. Unless otherwise noted, time between first tamoxifen administration and read-out was 10 weeks.

### Generation of mixed bone marrow chimeras

Bone marrow cells were harvested from WT CD45.1 or *Trac*^f/f^ E8_I_^Cre^ CD45.2 donors and depleted of T cell precursors using CD90.2 beads (Miltenyi) according to manufacturer’s instructions. An equal mix of 5×10^6^ total cells from WT CD45.1 and *Trac*^f/f^ E8_I_^Cre^ CD45.2 donors was intravenously injected into sub-lethally irradiated (6 Gy) *Rag1*^−/−^ hosts. Mice were analyzed 12-16 weeks after reconstitution.

### Single-cell TCR sequencing

Single cells were sorted using a FACS Aria into 96-well plates containing 5μL of lysis buffer (TCL buffer, Qiagen 1031576) supplemented with 1% β-mercaptoethanol) and frozen in −80°C prior to RT-PCR. RNA and RT-PCRs for TCRα and TCRβ were prepared as previously described (Dash et al., 2011). PCR products for TCRα and TCRβ were either subjected to Sanger sequencing or multiplexed with barcodes and subjected to MiSeq sequencing (Han et al., 2014) using True Seq Nano kit (Illumina). For Miseq data, Fastaq files were de-multiplexed and paired-end sequences assembled using PANDAseq (Masella et al., 2012) and FASTAX toolkit. Demultiplexed and collapsed reads were assigned to wells according to barcodes. Fasta files from both Sanger and Miseq sequences were aligned and analyzed on IMGT (imgt.org/HighV-QUEST) (Brochet et al., 2008). Cells with identical TCRβ CDR3 nucleotide sequences were considered as the same clones. Clonality was confirmed by sequencing TCRα of the expanded clones as assessed by TCRβ sequencing.

### Bulk RNAseq library preparation and analysis

Sorted cells (300-800 cells) were lysed in a guanidine thiocyanate buffer (TCL buffer, Qiagen) supplemented with 1% β-mercaptoethanol. RNA was isolated by solid-phase reversible immobilization bead cleanup using RNAClean XP beads (Agentcourt, A63987), reversibly transcribed, and amplified as described (Trombetta et al., 2014). Uniquely barcoded libraries were prepared using Nextera XT kit (Illumina) following manufacturer’s instructions. Sequencing was performed on an Illumina NextSeq550. Raw fastq files were pseudomapped against the mouse transcriptome (gencode M23) using the kallisto (v0.46) software (Bray et al., 2016). Transcript quantification was processed using the sleuth (v0.30) package for R (Pimentel et al., 2017). Shortly, we modeled batch effect and our experimental design using the sleuth_fit function and detected differentially expressed genes between all groups by the likelihood ratio test (lrt). When detecting significantly expressed genes between sample pairs, we used the wald-test function. Detected genes in the lrt and wald-test were used in downstream analysis if the minimum false discovery rate threshold of 0.05 and 1 log2 fold-change was reached. GSEA analysis was performed by using gene sets in gmt format and a full pre-ranked gene list by log2 fold-change between two groups as input for the fgsea package/R (Korotkevich et al., 2019). Gene ontology (GO) analysis was executed by comparing all detected genes as our background and the lists of differentially expressed genes against the biological processes gene sets by using the package for R topGO (Alexa and Rahnenfuhrer, 2019; Carlson M, 2019).

### Single cell RNAseq library preparation

IELs were sorted, counted for viability and immediately subjected to library preparation. The scRNA-seq and scTCR-seq libraries were prepared using the 10x Single Cell Immune Profiling Solution Kit, according to the manufacturer’s instructions at the Genomics core of Rockefeller University. The scRNA libraries were sequenced on an Illumina NextSeq550 to a minimum sequencing depth of 50,000 reads per cell using read lengths of 26 bp read 1, 8 bp i7 index, 98 bp read 2. The single-cell TCR libraries were sequenced on an Illumina NextSeq550 to a minimum sequencing depth of 5,000 reads per cell using read lengths of 150 bp read 1, 8 bp i7 index, 150 bp read 2.

### Data processing of single cell RNAseq and single cell TCRseq libraries

Raw fastq files derived from our RNA-seq libraries were processed with cellranger count (v3.1.0) using the 10x Genomics prebuilt mouse reference (v3.0.0 mm10). Our libraries were processed independently and merged into a single experiment at the analysis level using Seurat (v3.1.1) (Stuart et al., 2019). Quality control was performed by removing cells with high (> 5%) mitochondrial UMI content. Cells having more than 4000 or less than 200 genes were excluded from our analysis. TCR contigs and annotation were performed with the Cellranger vdj workflow from 10x Genomics and the prebuild mouse reference (v3.1.0 mm10). Paired TCR clonotypes were defined by the V, (D), J and CDR3 nucleotide composition for alpha and beta chains. Cells in which only one of the TCR sequences was recovered were excluded from the paired TCR clonal composition analysis. TCR clonotype sharing was assigned to cells expressing identical V, (D), J, and CDR3 nucleotide and amino acid sequences. Further processing and statistical analysis were performed using various R packages as described (RCoreTeam, 2019).

### Single cell RNAseq normalization and statistical analysis

The raw UMI counts were normalized by applying a regression model with negative binomial error distribution, available through the SCTransform function in the Seurat (v3.1.1) package (Hafemeister and Satija, 2019). The top 3000 variable genes were first used for dimensional reduction by PCA using the scaled data. The first 30 principal components were further used on the clustering algorithm and UMAP embedding for two dimensional visualization by using the Seurat workflow (Hafemeister and Satija, 2019; Stuart et al., 2019).

### Diffusion map and pseudotime analysis

To infer cell differentiation trajectories based on the expression data, we used a diffusion map algorithm adapted for sc-RNAseq analysis and implemented through the package destiny (Angerer et al., 2016). Normalized values were used as input for the Diffusion Map function. Cells were ordered based on the first diffusion component. To visualize lineage differentiation within our UMAP embedding and find differentially expressed genes over pseudotime, we used the slingshot package (Street et al., 2018).

### Statistical Analyses

Statistical analysis was carried out using GraphPad Prism v.8. Flow cytometry analysis was carried out using FlowJo software. Data in graphs show mean +/− SEM and p values <0.05 were considered significant. Repertoire diversity was analyzed by the Chao1 index and by Diversity 50 (D50). The Chao1 index (Chao, 1984), a measure of alpha diversity, was calculated using EstimateS software (Colwell et al., 2012). Diversity 50 (D50) was calculated on Excel as the fraction of dominant clones that account for the cumulative 50% of the total paired CDR3s identified in each UMAP cluster. CDR3 similarity (TCR sharing) was calculated using the Morisita-horn overlap index by using the divo package (Rempala and Seweryn, 2013). GraphPadPrism v.8 was used for graphs and Adobe Illustrator 2019 used to assemble and edit figures.

## Supplementary Table Titles

**Table S1. Top genes per UMAP cluster**

**Table S2. Clonotypes per UMAP cluster**

## Supplementary Figure Legends

**Figure S1.**
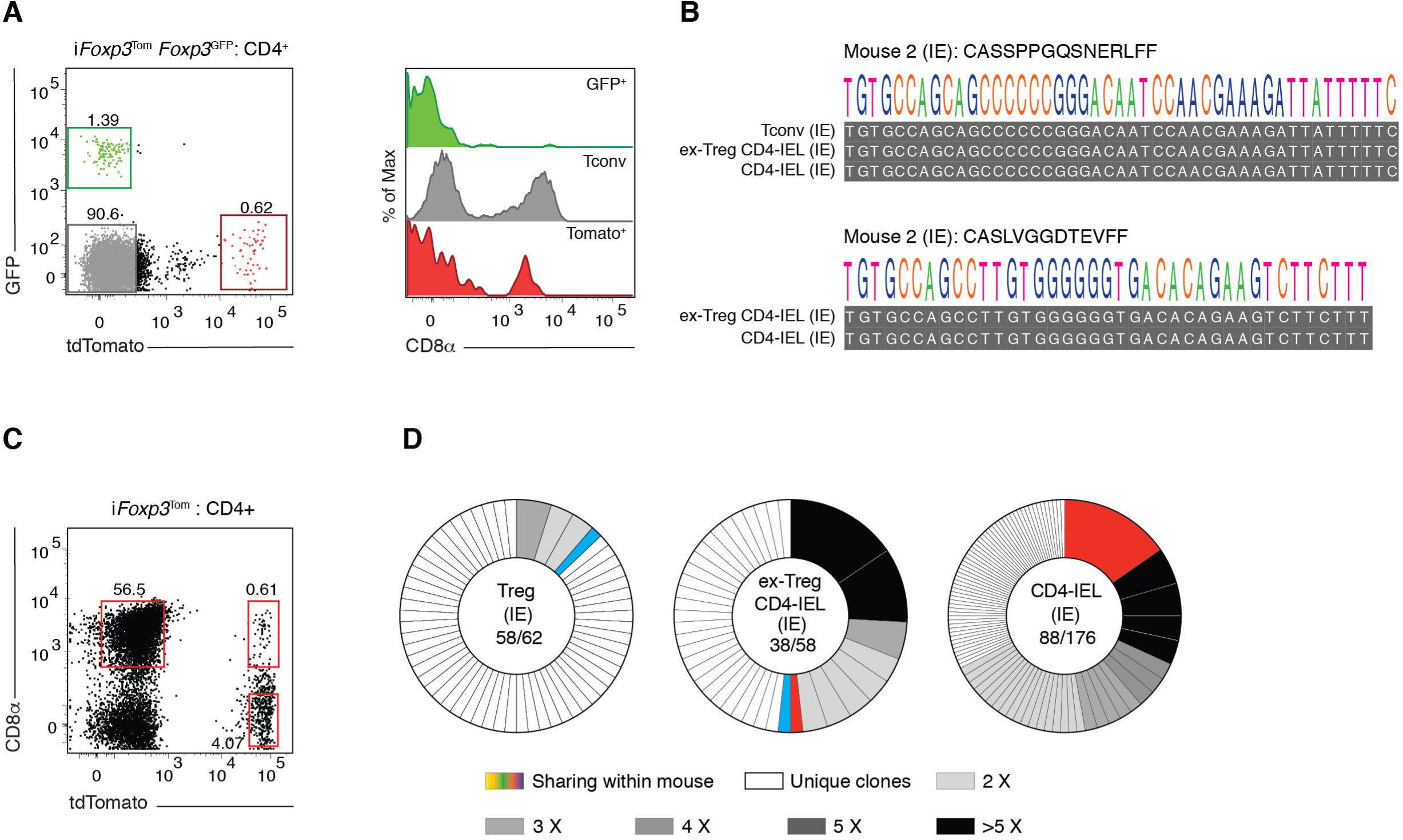
(related to Fig.1; CD4-IELs are clonally expanded with decreased TCR diversity). (**A, B**) *Foxp3* ^eGFP-Cre-ERT2^ x *Rosa26*^lsl-tdTomato^ x *Foxp3*^IRES-GFP^ (i*Foxp3*^Tom^ *Foxp3*^GFP^) mice were treated with tamoxifen for 10 weeks, and CD4^+^ T cells from mesenteric lymph nodes (mLN), lamina propria (LP) and intestinal epithelium (IE) were sorted as follows: CD4^+^ Conventional (Tconv; GFP^−^Tomato^−^CD8α^−^), regulatory T cells (Treg; GFP^+^ or Tomato^+^CD8α^−^), ex-Treg CD4-IEL (Tomato^+^CD8α^+^) and CD4-IEL (GFP^−^Tomato^−^ CD8α^+^). TCRβ (and TCRα of expanded TCRβ clones) were sequenced via the MiSeq platform. (**A**) Representative dot plot of GFP^+^ (green), tdTomato^+^ (red) and double-negative (grey) CD4^+^ T cells in the IE (left) and their corresponding CD8α expression (histogram, right). (**B**) Representative nucleotide sequence alignment of TCRβ CDR3 of 2 expanded shared clones. (**C**) Dot plot depicting CD8α and tdTomato expression by intraepithelial CD4^+^ T cells of an i*Foxp3*^*Tom*^ mouse. Sorted Tregs (Tomato^+^ CD8α^−^), ex-Treg CD4-IELs (Tomato^+^CD8α^+^) and CD4-IELs (Tomato^−^CD8α^+^) indicated by red boxes were sequenced by Sanger sequencing. (**D**) TCRβ clonal diversity of indicated IE populations. Each slice represents a distinct TCRβ CDR3. Colored clones represent sharing between populations. White slices represent unique clones and grey-scale slices represent expanded clones at indicated numbers. TCRα of expanded and/or shared clones were sequenced to confirm clonality.

**Figure S2.**
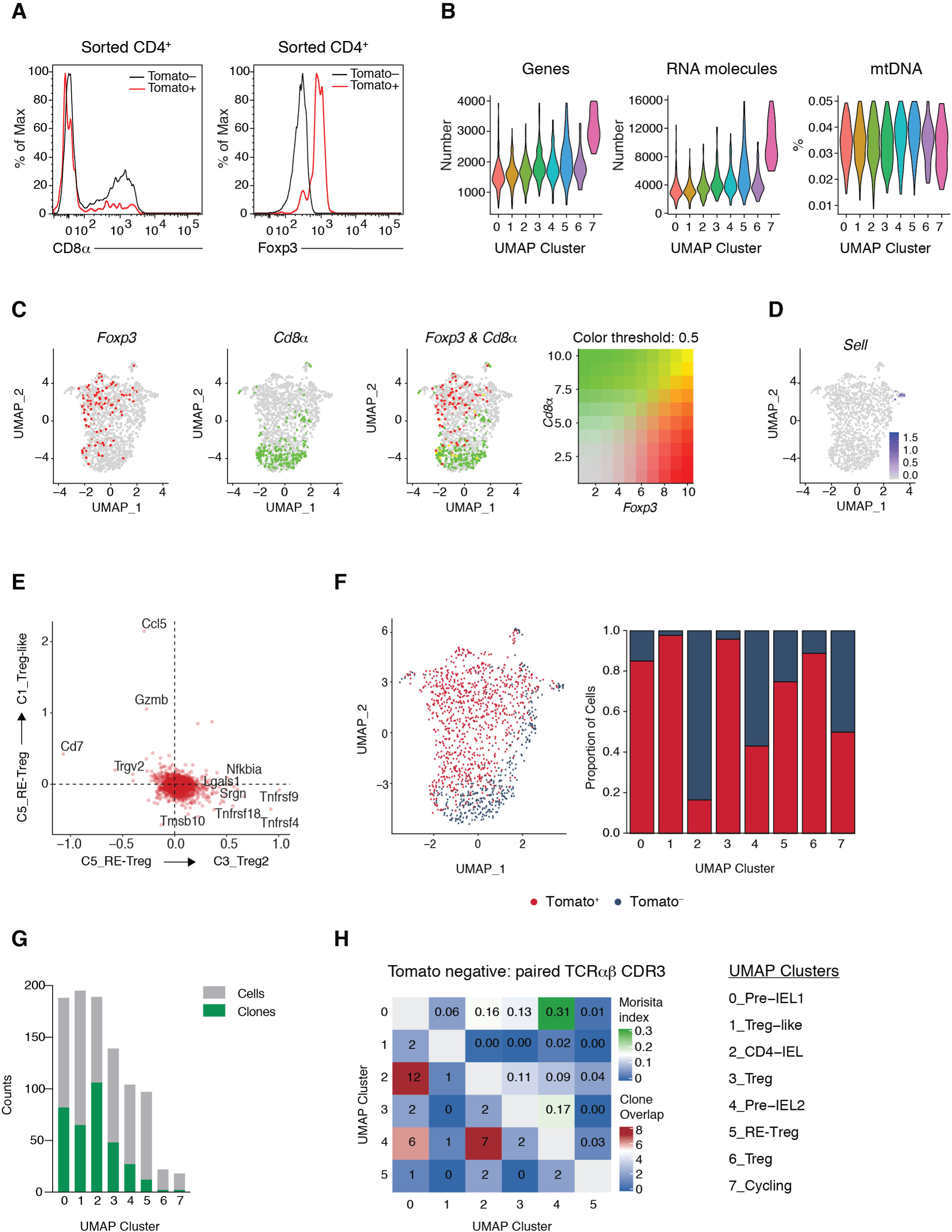
(related to Fig.2; Clonal distribution of intraepithelial CD4^+^ T cells follows the single-cell trajectories). (**A-F**) i*Foxp3*^Tom^ mice were treated with tamoxifen for 10 weeks, and Tomato^+^ (library 1) and Tomato^−^ (library 2) CD4^+^ T cells from the intestinal epithelium (IE) were sorted for scRNAseq using 10X Genomics platform. (**A**) Surface CD8α and intranuclear Foxp3 expression of sequenced CD4^+^ Tomato^+^ (red) and Tomato^−^ (black) cells. (**B**) Number of sequenced genes (left) and RNA molecules (middle) per cluster and percent of mitochondrial DNA (right) per UMAP cluster. (**C**) Expression levels of *Foxp3* (red), *Cd8a* (green), or both (yellow) by all analyzed cells. (**D**) Expression levels of *Sell* by analyzed cells. (**E**) 2D volcano plot comparing clusters 5(RE-Treg), 3 (Treg) and 1 (Treg-like). (**F**) Proportion of cells per UMAP cluster from library 1 (Tomato^+^, red) and library 2 (Tomato^−^, blue). (**G**) Total number of cells with paired αβTCR sequences (grey) and total number of clones (green) within each UMAP cluster. (**H**) Normalized Morisita index (top right) and number of shared clones (bottom left) of paired αβTCR per UMAP cluster among tomato^−^ cells.

**Figure S3.**
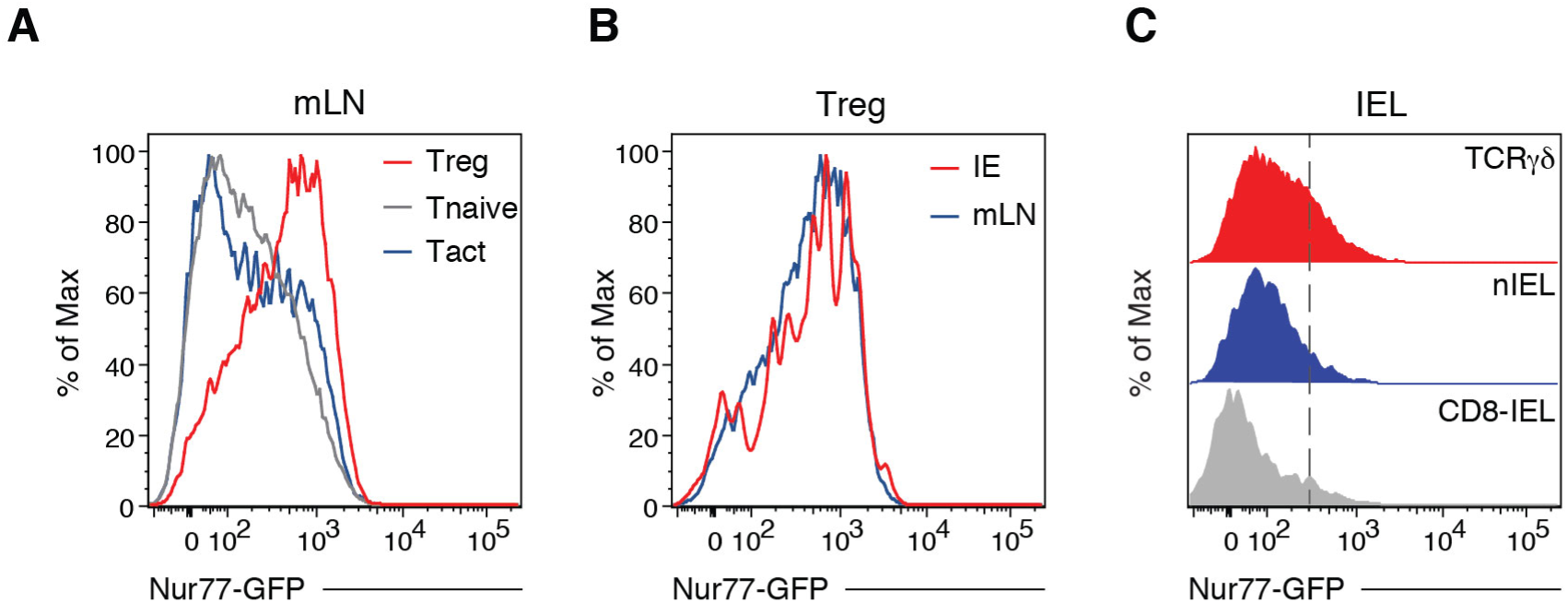
(related to Fig.3; Intraepithelial CD4^+^ T cell subsets show an inverse correlation between TCR signaling and IEL program). (**A-C**) *Nur77*-GFP expression levels by indicated cell types from *Nur77*^GFP^ *Foxp3*^RFP^ double-reporter mice. (**A**) *Nur77*^GFP^ expression among CD4^+^ Foxp3^+^ regulatory T cells (Treg, red), CD62L^high^CD44^low^ naïve T cells (Tnaïve, grey), and CD62L^low^CD44^high^ Foxp3^−^ activated T cells (Tact, blue) in the mesenteric lymph nodes (mLN). (**B**) *Nur77*^GFP^ expression among Foxp3^+^ Tregs in the intestinal epithelium (IE, red) and mLN (blue). (**C**) *Nur77*^GFP^ expression among TCRγδ-IELs (red), CD8αα^+^ CD8β^−^CD4^−^TCRαβ^+^ natural IELs (nIEL, blue) and CD8αα^+^ CD8β^+^ TCRαβ^+^ IELs (CD8-IEL, grey) in the small intestine epithelium.

**Figure S4.**
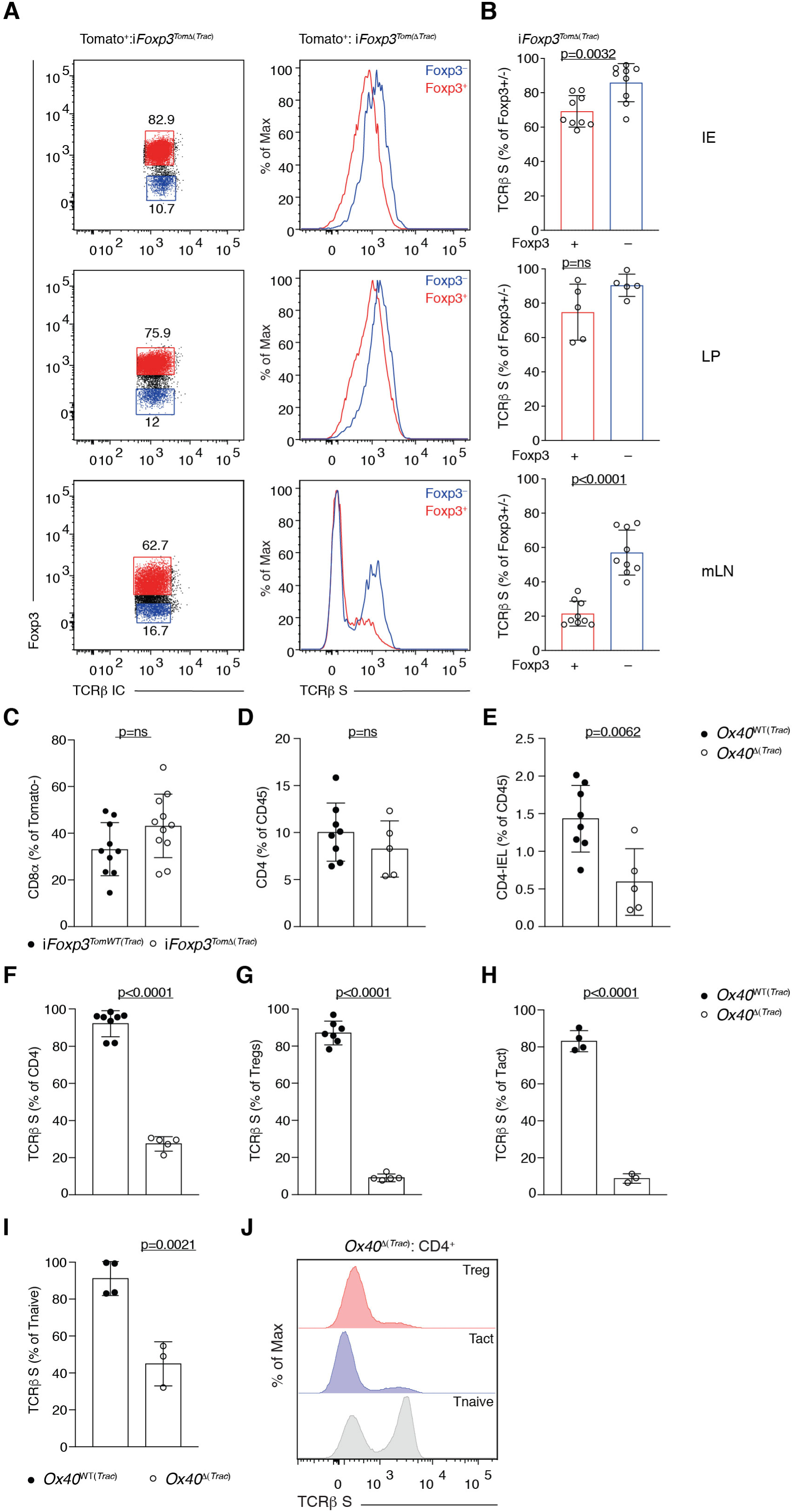
(related to Fig.4; TCR signaling is required for CD4-IEL differentiation). (**A-C**) Flow cytometry analysis of i*Foxp3*^*Tom*WT*(Trac)*^ (*Trac*^+/+^ i*Foxp3*^Tom^) or i*Foxp3*^*Tom*Δ(*Trac)*^ (*Trac*^f/f^ i*Foxp3*^Tom^) mice 8-12 weeks after tamoxifen administration. (**A**) Representative dot plots (left) for intracellular TCRβ and Foxp3, or histograms (right) of surface TCRβ expression among tomato^+^ Foxp3^+^ Tregs (red) and tomato^+^ Foxp3^−^ CD4^+^ T cells (blue) in the intestinal epithelium (IE, top), lamina propria (LP, middle) and mesenteric lymph nodes (mLN, bottom). (**B**) Frequencies of surface TCRβ-expressing cells among Foxp3^+^ or Foxp3^−^ among tomato^+^ CD4^+^ T cells from i*Foxp3*^*Tom*Δ(*Trac)*^ IE (top), LP (middle) or mLN (bottom). (**C**) Frequency of CD8α-expressing CD4^+^ T (CD4-IEL) cells among tomato^−^ cells in the IE of i*Foxp3*^*Tom*WT(*Tra*c)^ or i*Foxp3*^*Tom*Δ(*Trac)*^ mice. (**D-J**) Flow cytometry analysis of cells isolated from the IE and mLN of 9-12 week-old OX40^WT(*Trac)*^ (*Trac*^+/+^ OX40Cre^+/−^ or *Trac*^f/f^ OX40Cre^−/−^) or OX40^Δ(*Trac)*^ (*Trac*^f/f^ OX40Cre^+/−^) mice. (**D, E**) Frequencies of total CD4^+^ T cells (**D**) and CD4-IELs (CD8α^+^TL-Tetramer^+^) (**E**) among total CD45^+^ cells in the IE. (**F-J**) Frequencies of surface TCRβ-expressing cells among CD4^+^ T cells (**F**), Tregs (**G**), CD4^+^ activated (Tact, Foxp3^−^CD44^high^CD62L^low^) (**H**) and CD4^+^ Naïve (Tnaïve, Foxp3^−^CD44^low^CD62L^high^) (**I**) cells in the mLN. (**J**) Histogram of surface TCRβ expression among Tregs (red), Tact (blue) and Tnaïve (grey) CD4^+^ T cells in the mLN. Data are expressed as mean +/− SEM of individual mice (n=5-11). Significant p values as indicated [student’s t test (**B-I**)].

**Figure S5.**
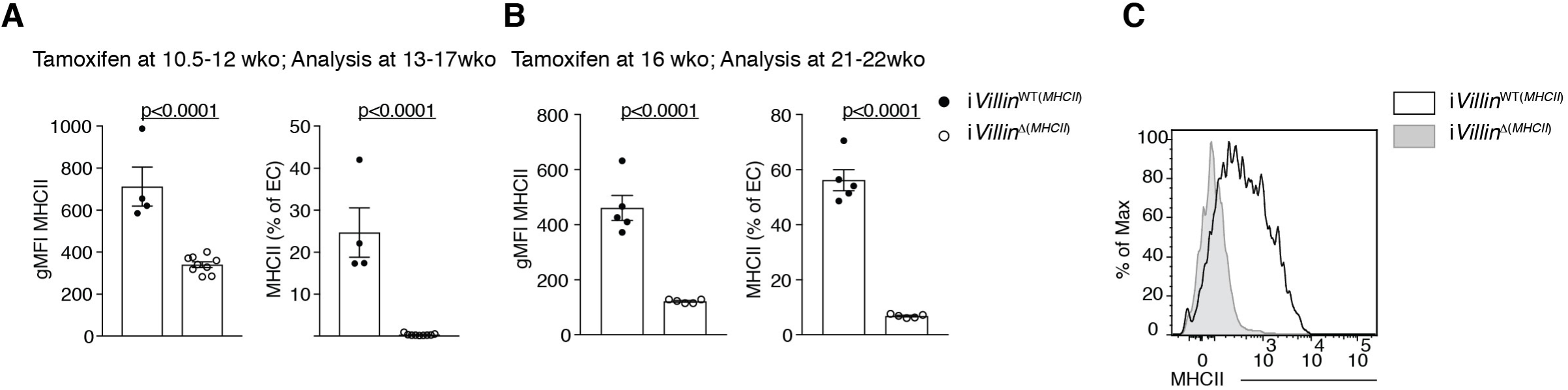
(related to Fig.5; MHCII expression by epithelial cells is required for CD4-IEL conversion). (**A-C**) Flow cytometry analysis of the intestinal epithelium (IE) of i*Villin*^WT(*MHCII)*^ (*H2-Ab1*^+/+^ Villin^CreERT2+/–^ or *H2-Ab1*^f/f^ Villin^CreERT2–/–^) or i*Villin*^Δ(*MHCII)*^ (*H2-Ab1*^f/f^ Villin^CreERT2+/+^) mice after tamoxifen administration. Geometric mean fluorescence intensity (gMFI) (left) and frequency (right) of MHCII expression by epithelial cells of 13-17 week-old mice 3-5 weeks after tamoxifen administration (**A**) and of 21-22 week-old mice 5 weeks after tamoxifen administration (**B**). (**C**) Representative histogram of MHCII expression by EpCAM^+^ epithelial cells in 20 week-old i*Villin*^WT(*MHCII)*^ (white) or i*Villin*^Δ(*MHCII)*^ (grey) mice 4 weeks after tamoxifen administration. Data are expressed as mean +/− SEM of individual mice (n=5-14). Significant p values as indicated [student’s t test (**A, B**)].

**Figure S6.**
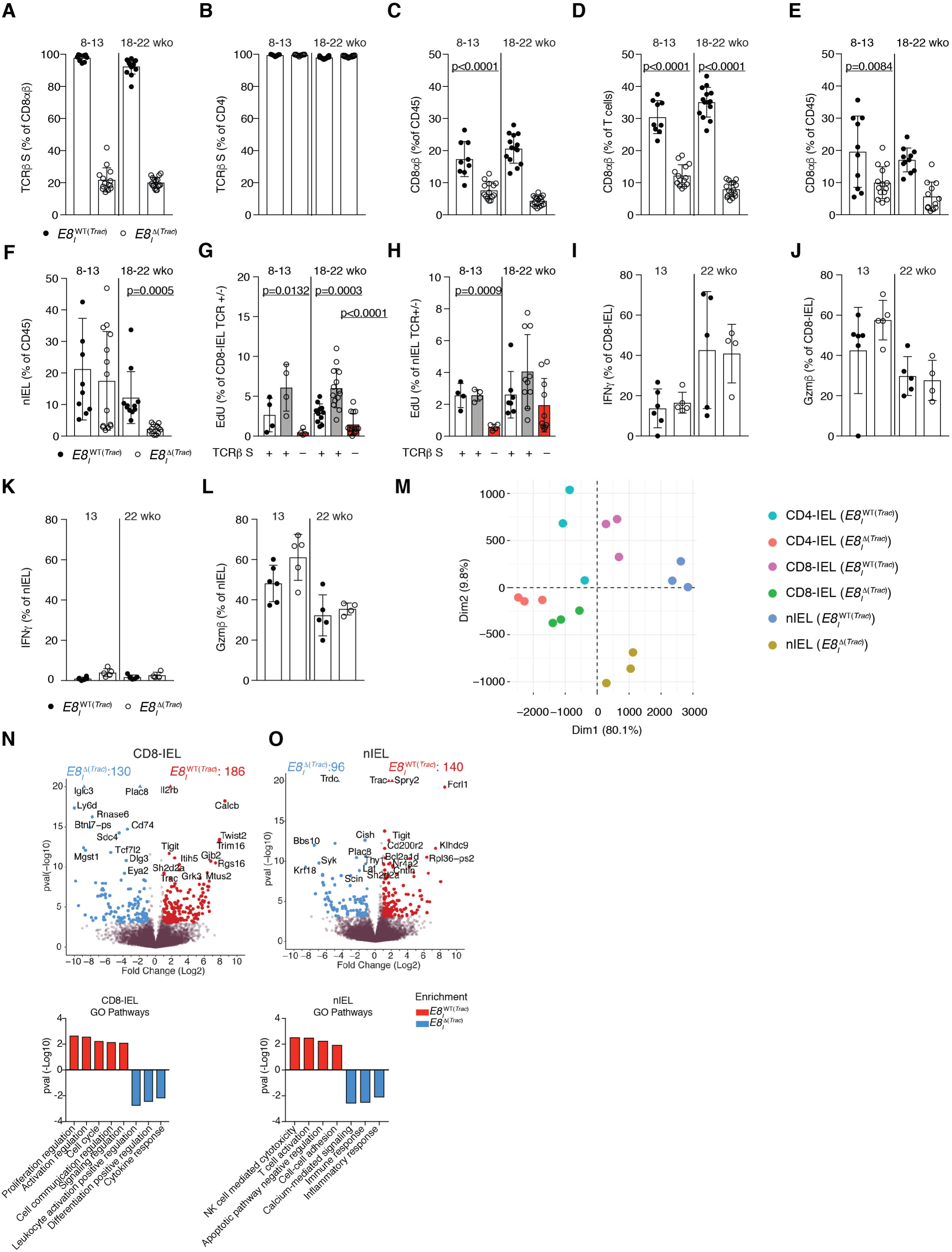
(related to Fig.6; TCR signaling is not essential for CD4-IEL maintenance). (**A-L)** Flow cytometry analysis of 8-22 week-old E8_I_^WT(*Trac)*^ (*Trac*^+/+^ E8_I_Cre^+^ or *Trac*^+/+^ E8_I_Cre^−^ or *Trac*^f/f^ E8_I_Cre^−^) or E8 _I_^Δ(*Trac)*^ (*Trac*^f/f^ E8_I_Cre^+^) mice, grouped by age as indicated. Frequencies of surface TCRβ-expressing cells among CD8αβ cells (**A**) and CD4 (**B**) T cells in the mesenteric lymph nodes (mLN). Frequency of CD8αβ T cells among total CD45^+^ cells (**C**) and intracellular TCRβ-expressing T cells (**D**) in the mLN. Frequencies of CD8αβ (**E**) and natural IELs (nIEL, CD4^−^CD8αα^+^CD8β^−^TL-Tetramer^+^) (**F**) among CD45^+^ cells in the intestinal epithelium (IE). (**G, H**) Proliferation (EdU incorporation) of CD8-IELs (**G**) or nIELs (**H**) with or without surface TCRβ expression after EdU injection 16 and 4 hours prior to analysis. (**I-L**) Frequencies of IFNγ (**I, K**) and Gzmb (**J, L**) production upon PMA/Ionomycin *ex-vivo* stimulation among CD8-IELs (**I, J**) and nIELs (**K, L**). (**M-O**) Bulk RNA-sequencing was performed on TCR^+^ CD4-IELs, CD8-IELs and nIELs from E8_I_^WT(*Trac)*^ and TCR^−^ CD4-IELs from E8_I_^Δ(*Trac)*^ mice. (**M**) Principal component analysis of indicated cell populations. (**N**,**O**) Volcano plots of differentially expressed genes between indicated populations (p<0.05, in color) (top), and selected differentially-enriched gene ontology (GO) pathways between them (bottom). Data are expressed as mean +/− SEM of individual mice. (n=4-14). Significant p values as indicated [student’s t test (**A-F, I-L**) or one-way ANOVA and Bonferroni (**G, H**)]. N=10-16 (**A-F**), n=4-14 (**G, H**) and n=4-6 (**I-L**). Sequencing data is n=3 mice per group (**M-O**).

