## Supplementary material for "T cell receptor is required for differentiation but not maintenance of intestinal intraepithelial lymphocytes": Table S3

### Table S3. Differentially expressed genes between $E8_I^{WT(\textit{Trac})}$ and $E8I^{\Delta(\textit{Trac})}$ CD4-IELs

| Overexpressed in $E8_I^{WT(\textit{Trac})}$ | | | Overexpressed in $E8_I^{\Delta(\textit{Trac})}$ | | |
|---|---|---|---|---|---|
| **gene** | **ave_logFC** | **p_val_adj** | **gene** | **ave_logFC** | **p_val_adj** |
| Cd83 | 8.577585 | 2.93E-10 | Gm21762 | -10.5303 | 1.67E-20 |
| Klrb1a | 6.074966 | 8.77E-10 | Klk1 | -9.84137 | 1.74E-13 |
| Klrb1c | 8.030817 | 3.8E-09 | Btla | -7.32358 | 5.65E-08 |
| Klre1 | 4.849967 | 4.02E-09 | Bcl11a | -6.9866 | 3.84E-06 |
| Spry2 | 5.75853 | 4.02E-09 | Cox6a2 | -9.01962 | 5.51E-06 |
| Lag3 | 2.264442 | 7.29E-09 | Fcrla | -7.98591 | 8.32E-06 |
| Gm19590 | 7.492748 | 2.45E-06 | Gm17021 | -7.08457 | 3.41E-05 |
| Gm16299 | 7.230879 | 3.84E-06 | Cct6a | -1.04941 | 3.97E-05 |
| Bcl2a1b | 1.348215 | 4.8E-06 | Dnase1l3 | -5.57644 | 0.000122 |
| Zdhhc15 | 3.332176 | 5.51E-06 | Alkbh3os1 | -6.74636 | 0.000178 |
| Ky | 4.523866 | 1.38E-05 | Zfp607b | -3.9259 | 0.000315 |
| Gm4956 | 4.323781 | 2.86E-05 | Gm43375 | -6.5652 | 0.00034 |
| Cnrip1 | 6.095595 | 5.15E-05 | Gm38214 | -6.29071 | 0.000786 |
| Gins1 | 5.599377 | 8.68E-05 | Gm43196 | -3.80262 | 0.001025 |
| Kazald1 | 6.476025 | 9.52E-05 | Olfr99 | -5.29606 | 0.001253 |
| Cd200r1 | 1.51688 | 0.000134 | Sell | -6.53934 | 0.001317 |
| Spin2c | 6.760562 | 0.000135 | Gm49179 | -7.03964 | 0.001445 |
| Gm1821 | 1.658691 | 0.000147 | Dntt | -8.32337 | 0.001759 |
| Zfp932 | 2.325318 | 0.000204 | Olfr528-ps1 | -5.79965 | 0.001884 |
| Nrp2 | 3.161478 | 0.000387 | Tlr6 | -4.83628 | 0.002789 |
| Gm17928 | 3.056256 | 0.000687 | Gm37321 | -5.22711 | 0.002818 |
| Cyp2j9 | 6.412093 | 0.000786 | Lyst | -2.68811 | 0.003276 |
| Bcl2a1d | 1.583238 | 0.001253 | Gm50221 | -5.98038 | 0.003785 |
| Tspan4 | 5.606894 | 0.001253 | Fbxl13 | -6.66278 | 0.003851 |
| Gm15996 | 3.612176 | 0.001445 | Gm26575 | -5.36307 | 0.004354 |
| Gm42611 | 2.160161 | 0.001884 | 1500002F19Rik | -5.90029 | 0.004702 |
| Hist1h2ap | 2.174963 | 0.002789 | Trav9-2 | -2.84198 | 0.005862 |
| Tnfrsf9 | 4.01641 | 0.002789 | Siglech | -5.70372 | 0.006438 |
| A930001A20Rik | 6.932589 | 0.003114 | Flt3 | -3.48482 | 0.007187 |
| Tmem30a | 1.048406 | 0.00345 | Gm33882 | -5.21009 | 0.007605 |
| Rorc | 5.216056 | 0.00345 | Gm44958 | -6.21954 | 0.008696 |
| Pla2g4b | 5.985127 | 0.00345 | Wdr13 | -2.58232 | 0.009802 |
| Golph3 | 1.529466 | 0.003785 | Mynn | -1.66741 | 0.01207 |
| Gm14435 | 3.132717 | 0.003795 | Cfap97 | -1.59325 | 0.013068 |
| Zfp160 | 2.240651 | 0.003997 | Gm37357 | -1.68319 | 0.013597 |
| Egr3 | 5.691184 | 0.004354 | Ppfia4 | -7.76848 | 0.016735 |
| Zdhhc3 | 1.780378 | 0.004517 | Mtfp1 | -4.89168 | 0.017021 |
| Gm11802 | 5.439506 | 0.004591 | Fam220a | -1.48648 | 0.017486 |
| Trav7d-5 | 6.150892 | 0.004702 | Gm48939 | -4.25497 | 0.021422 |
| Trav7n-5 | 6.150892 | 0.004702 | Gm45495 | -3.3197 | 0.023629 |
| Nfkbib | 1.538276 | 0.005151 | Plaur | -4.72063 | 0.032266 |
| Stmn1 | 2.374753 | 0.006036 | Gm28373 | -4.53106 | 0.034206 |
| Setmar | 4.208802 | 0.006036 | Cd300c | -7.77426 | 0.034914 |
| Tsga10 | 4.828529 | 0.006226 | Plag1 | -5.82179 | 0.034914 |
| Il3ra | 4.934256 | 0.006226 | Appbp2 | -1.16342 | 0.034914 |
| Fam174b | 5.608441 | 0.006704 | 9030407P20Rik | -3.58214 | 0.03579 |
| Cks2 | 1.710192 | 0.00711 | Plce1 | -2.54427 | 0.036514 |
| Plek | 3.135804 | 0.008186 | Gm16060 | -3.98283 | 0.037372 |
| Gm47015 | 1.492324 | 0.008696 | Adcy3 | -2.10636 | 0.039316 |
| Enkd1 | 3.061886 | 0.008868 | Lrp8 | -3.96959 | 0.039567 |
| Gm16572 | 5.696436 | 0.009159 | Rlf | -1.53071 | 0.040456 |
| Creb3l1 | 5.436078 | 0.00985 | Rbm33 | -1.44965 | 0.040456 |
| Osgin1 | 1.476867 | 0.010278 | Ltb4r1 | -4.5426 | 0.043707 |
| Cenpf | 3.138301 | 0.010639 | Trav13-1 | -2.70602 | 0.045393 |
| Zfp458 | 2.6881 | 0.010772 | Trav10 | -2.52101 | 0.045947 |
| Pnpo | 1.792949 | 0.012193 | | | |
| Clec16a | 2.552224 | 0.012193 | | | |
| Ska1 | 5.192346 | 0.012193 | | | |
| Bcap29 | 1.224712 | 0.01458 | | | |
| Rpl36-ps2 | 4.058048 | 0.01458 | | | |
| Gem | 5.760524 | 0.014852 | | | |
| Gm11423 | 2.545579 | 0.015383 | | | |

| | | |
|---|---|---|
| Vps54 | 1.181797 | 0.017021 |
| Itih5 | 3.820922 | 0.017109 |
| Mlkl | 4.21895 | 0.017486 |
| Rictor | 1.43574 | 0.018067 |
| 6330403K07Rik | 3.216177 | 0.0202 |
| Rida | 1.775842 | 0.021422 |
| Aoah | 4.281722 | 0.021745 |
| Mbnl3 | 1.335676 | 0.022484 |
| Ephx1 | 2.189419 | 0.024468 |
| Txnl4b | 1.785039 | 0.027541 |
| Gm25363 | 4.259935 | 0.029876 |
| Ttc30a1 | 6.915919 | 0.03179 |
| Glp1r | 1.449691 | 0.032266 |
| Ciart | 4.260166 | 0.032266 |
| As3mt | 2.00779 | 0.032315 |
| Gm36262 | 4.862997 | 0.032958 |
| Igflr1 | 1.165425 | 0.033625 |
| Gm47830 | 4.834154 | 0.034206 |
| Gm26510 | 1.513678 | 0.036514 |
| Nat14 | 4.424568 | 0.037372 |
| Zfp212 | 1.779624 | 0.038153 |
| Nup133 | 2.660052 | 0.039147 |
| Ucp2 | 1.135401 | 0.040002 |
| Kat6a | 1.507153 | 0.041213 |
| Gm48735 | 1.834047 | 0.041332 |
| Penk | 3.417682 | 0.041658 |
| Retreg3 | 1.891107 | 0.042317 |
| Pgm1 | 1.312999 | 0.043707 |
| Tacc2 | 1.768766 | 0.043707 |
| Cog7 | 1.98717 | 0.044089 |
| Ylpm1 | 1.194531 | 0.044847 |
| Lrrc20 | 1.862899 | 0.044847 |
| Gm45110 | 2.411711 | 0.045947 |
| Egr1 | 5.215026 | 0.04765 |
