## Supplementary material for "T cell receptor is required for differentiation but not maintenance of intestinal intraepithelial lymphocytes": Table S4

### Table S4. Differentially expressed genes between E8I$^{WT(\textit{Trac})}$ and E8I$^{\Delta(\textit{Trac})}$ CD8-IELs

| Overexpressed in E8I$^{WT(\textit{Trac})}$ | | | Overexpressed in E8I$^{\Delta(\textit{Trac}}$ | | |
|---|---|---|---|---|---|
| **gene** | **ave_logFC** | **p_val_adj** | **gene** | **ave_logFC** | **p_val_adj** |
| Il2rb | 1.838553 | 6.58E-18 | Plac8 | -1.89668 | 1.78E-26 |
| Calcb | 8.540629 | 3.06E-15 | Iglc3 | -8.77103 | 5.07E-17 |
| Twist2 | 7.857668 | 7.89E-11 | Ly6d | -9.98308 | 1.9E-14 |
| Trim16 | 7.818193 | 1.3E-10 | Rnase6 | -7.78589 | 1.99E-13 |
| Tigit | 1.669783 | 2.94E-09 | Btnl7-ps | -8.08394 | 3.67E-12 |
| Tsga10 | 2.336488 | 9.2E-09 | Cd74 | -3.47087 | 5.08E-12 |
| Gjb2 | 6.788909 | 2E-08 | Sdc4 | -4.4874 | 1.3E-11 |
| Rgs16 | 7.370943 | 3.09E-08 | Ppp1r14d | -8.84606 | 7.01E-10 |
| Itih5 | 2.927409 | 4.68E-08 | Mgst1 | -8.59324 | 1.35E-09 |
| Grk3 | 2.914981 | 2.08E-07 | Tcf7l2 | -5.52242 | 2.29E-09 |
| Mtus2 | 6.363309 | 4.5E-07 | Dlg3 | -3.61153 | 1.75E-08 |
| Sh2d2a | 1.038083 | 4.79E-07 | Eya2 | -3.81918 | 4.91E-07 |
| Rpl36-ps2 | 5.473854 | 1.93E-06 | Heatr5a | -1.45863 | 2.77E-06 |
| Spry2 | 1.78517 | 2.08E-06 | Slc51a | -4.2348 | 2.89E-06 |
| Phtf1os | 6.594028 | 3.61E-06 | Gzmk | -1.25234 | 3.41E-06 |
| 1700047G07Rik | 2.376492 | 6.52E-06 | Clca4b | -9.82448 | 3.61E-06 |
| Nr4a1 | 5.926869 | 7.91E-06 | Smim22 | -7.36771 | 5.48E-06 |
| Nusap1 | 5.378206 | 7.91E-06 | Cds1 | -6.52016 | 6.2E-06 |
| Gm45716 | 2.275984 | 1.27E-05 | Cdc42ep4 | -6.15667 | 6.2E-06 |
| Cdkn3 | 6.381035 | 1.32E-05 | Gprin3 | -6.52167 | 2.09E-05 |
| Wnt10a | 6.549708 | 1.35E-05 | Thy1 | -1.40797 | 2.09E-05 |
| Plek | 1.972456 | 1.35E-05 | Atp1b1 | -3.07396 | 2.18E-05 |
| Ephx1 | 2.602742 | 2.09E-05 | Glipr2 | -3.34186 | 2.3E-05 |
| Ckap2l | 5.762481 | 2.09E-05 | Kcnj8 | -6.59678 | 2.79E-05 |
| Serp2 | 4.215937 | 2.63E-05 | Ptprf | -6.41495 | 2.9E-05 |
| Dram1 | 6.451004 | 2.9E-05 | Ifit1bl1 | -1.94358 | 4.31E-05 |
| Crem | 2.716409 | 3.27E-05 | Prss30 | -6.19576 | 7.83E-05 |
| 2900026A02Rik | 1.467805 | 3.27E-05 | Gpa33 | -6.5909 | 9.16E-05 |
| Birc5 | 3.068787 | 3.44E-05 | 4930447K03Rik | -6.24899 | 9.16E-05 |
| 6720464F23Rik | 1.830093 | 4.56E-05 | Gm39091 | -5.62813 | 0.00011 |
| Klrb1a | 4.024957 | 5.03E-05 | Pak6 | -6.75568 | 0.000113 |
| Nrn1 | 5.842732 | 5.29E-05 | Rnf186 | -7.1091 | 0.000119 |
| Ptpn6 | 1.302889 | 0.000113 | Btbd16 | -2.15688 | 0.000146 |
| Cdc6 | 5.69186 | 0.000114 | Trim31 | -6.1468 | 0.000181 |
| 9530050K03Rik | 4.68472 | 0.000119 | Crb3 | -2.5452 | 0.000243 |
| Ska1 | 5.539225 | 0.000128 | Dkkl1 | -1.8539 | 0.000244 |
| Cpa2 | 6.142104 | 0.000151 | Gtf3c4 | -2.53712 | 0.00033 |
| Tox | 1.322971 | 0.000202 | Il7r | -2.17182 | 0.000428 |
| Gm43636 | 3.863844 | 0.000225 | Soat2 | -1.02958 | 0.000455 |
| Iqsec1 | 1.396754 | 0.000232 | Lypd8 | -10.6959 | 0.000476 |
| Gm156 | 3.508176 | 0.000239 | Klf3 | -2.15986 | 0.000542 |
| Gm48182 | 5.344042 | 0.000274 | H2-Ab1 | -3.23751 | 0.000642 |
| Igflr1 | 1.194761 | 0.00033 | Phf11b | -2.10252 | 0.000652 |
| Gm38190 | 1.094031 | 0.000365 | Pou4f1 | -3.14446 | 0.000669 |
| Gm30790 | 5.7876 | 0.000388 | Gdpd1 | -5.81388 | 0.000727 |
| Mnd1 | 5.976298 | 0.000415 | Cdr2 | -2.52604 | 0.000759 |
| Pdzd2 | 2.764092 | 0.000423 | Lta | -2.9727 | 0.000775 |
| Bcl2l11 | 1.152136 | 0.000444 | Gm36043 | -2.6426 | 0.001065 |
| Tnfsf4 | 5.938302 | 0.00049 | Tm4sf5 | -3.21862 | 0.001082 |
| Hist1h2ap | 1.655867 | 0.000535 | Slc5a1 | -9.46691 | 0.001115 |
| Nepro | 2.364155 | 0.000554 | F2rl2 | -2.12807 | 0.001198 |
| Zfp935 | 6.530183 | 0.000572 | Lgr4 | -4.04175 | 0.00123 |
| Gm44901 | 1.768357 | 0.000577 | Ptafr | -3.26735 | 0.00123 |
| Slc35g2 | 5.537121 | 0.000642 | Cd24a | -5.67781 | 0.001467 |
| S100a1 | 1.906517 | 0.00071 | Mthfr | -5.57881 | 0.001816 |
| Zfp318 | 2.623999 | 0.000727 | S100g | -4.58647 | 0.001886 |
| Rnf19a | 2.466756 | 0.000731 | App | -2.26836 | 0.001928 |
| Cd101 | 1.255209 | 0.000804 | Gm2237 | -2.79401 | 0.001977 |
| 4833445I07Rik | 5.727192 | 0.000969 | Nfatc2 | -1.41669 | 0.00229 |
| Dusp5 | 1.972161 | 0.001065 | Slc4a10 | -3.39131 | 0.0036 |
| M1ap | 3.148858 | 0.001065 | Mansc4 | -5.44433 | 0.004484 |
| Unc13a | 4.269183 | 0.00123 | Src | -5.01571 | 0.00449 |

| | | |
|---|---|---|
| Bcl2a1b | 1.37234 | 0.001232 |
| Gm12036 | 2.784046 | 0.001346 |
| Gm20689 | 6.455044 | 0.00143 |
| Cnih3 | 5.229141 | 0.00163 |
| Lag3 | 2.059385 | 0.00173 |
| Cib2 | 3.854989 | 0.001848 |
| Trav9d-3 | 2.29694 | 0.001856 |
| Trav9n-3 | 2.29694 | 0.001856 |
| Cep85l | 2.013048 | 0.001856 |
| Gm14377 | 2.262636 | 0.001856 |
| Ceacam-ps1 | 5.748506 | 0.001886 |
| Kif11 | 2.632826 | 0.001886 |
| Tnfrsf9 | 3.015079 | 0.001886 |
| Endod1 | 2.086059 | 0.001886 |
| Gm17928 | 2.721317 | 0.001886 |
| Gm44860 | 4.017727 | 0.001966 |
| Pabpc4 | 1.348914 | 0.00198 |
| Zfp36l1 | 1.330951 | 0.002061 |
| Ypel2 | 2.061505 | 0.002138 |
| Dars2 | 2.221242 | 0.002253 |
| Gmpr | 4.610124 | 0.002282 |
| C1qbp | 1.010959 | 0.002475 |
| Nrcam | 4.278282 | 0.0036 |
| Gm19590 | 2.897056 | 0.003952 |
| Trav4d-4 | 6.622715 | 0.004201 |
| Trav4n-4 | 6.622715 | 0.004201 |
| Bcl2l15 | 2.359205 | 0.00449 |
| Gm15287 | 3.360704 | 0.004735 |
| Il6st | 1.170593 | 0.004781 |
| Stat4 | 1.089641 | 0.004801 |
| Sulf2 | 2.2495 | 0.004921 |
| Gm44752 | 3.64351 | 0.005005 |
| Ccdc102a | 1.601084 | 0.00514 |
| Gm16638 | 2.277832 | 0.005644 |
| Galnt4 | 2.330559 | 0.005662 |
| Helz | 1.22547 | 0.005887 |
| Tctex1d1 | 4.153338 | 0.005903 |
| Gm816 | 4.302373 | 0.005903 |
| A530041M06Rik | 1.451263 | 0.005993 |
| Senp8 | 5.329353 | 0.006425 |
| Gm16575 | 4.086798 | 0.006452 |
| Ighv6-4 | 5.320397 | 0.007003 |
| Fam83d | 5.76987 | 0.007041 |
| Cd200r2 | 2.3063 | 0.007041 |
| Gm48904 | 2.976209 | 0.007674 |
| BB557941 | 5.118324 | 0.007736 |
| Wnk3 | 4.884693 | 0.007752 |
| Cd200r1 | 1.846769 | 0.00815 |
| Trbj1-1 | 2.12548 | 0.00858 |
| Got2-ps1 | 5.089218 | 0.008827 |
| Trav16d-dv11 | 5.299982 | 0.008938 |
| Gm14021 | 5.048681 | 0.009171 |
| Xlr3b | 4.612688 | 0.009539 |
| Ptrhd1 | 1.903929 | 0.009642 |
| Ehmt1 | 1.514238 | 0.010158 |
| Gm45406 | 4.974128 | 0.012009 |
| Ltf | 4.987067 | 0.0123 |
| Tnfsf8 | 2.271699 | 0.013892 |
| Arap3 | 1.686096 | 0.014451 |
| Bcl2a1d | 1.772999 | 0.01465 |
| Lilr4b | 1.425025 | 0.014869 |
| E230029C05Rik | 2.421245 | 0.015039 |
| Gm16741 | 3.995124 | 0.016117 |
| Glp1r | 1.029124 | 0.016762 |
| Capn3 | 1.017118 | 0.018631 |
| Cela1 | 2.111948 | 0.018868 |
| Pecam1 | 1.200806 | 0.018949 |

| | | |
|---|---|---|
| Ski | -1.5631 | 0.004634 |
| Tmem86a | -4.87196 | 0.005005 |
| Taf4 | -2.27375 | 0.006452 |
| Cish | -1.14121 | 0.006919 |
| Trpv2 | -1.31028 | 0.007024 |
| Myo7b | -5.99949 | 0.007253 |
| Mirt1 | -1.73149 | 0.007375 |
| Tnfaip1 | -2.0164 | 0.007449 |
| Cox6a2 | -5.60526 | 0.007597 |
| Arhgef11 | -6.1055 | 0.007849 |
| Rbm47 | -2.11455 | 0.007981 |
| Tmem97 | -1.09369 | 0.00858 |
| Rogdi | -2.46868 | 0.008938 |
| Ces2e | -8.64236 | 0.009581 |
| Kdelr1 | -1.6114 | 0.010074 |
| Ap1m2 | -3.36303 | 0.011595 |
| 5430431A17Rik | -1.14449 | 0.012706 |
| Lactb2 | -1.4014 | 0.014797 |
| Trav13d-3 | -5.12508 | 0.014838 |
| Cmbl | -6.38465 | 0.015846 |
| Cdkn2b | -4.55511 | 0.016762 |
| Mttp | -3.84505 | 0.016762 |
| Aoc1 | -7.88392 | 0.017424 |
| Tagln2 | -1.48519 | 0.01779 |
| Cxcr3 | -1.52725 | 0.018036 |
| Pde11a | -2.13368 | 0.018923 |

| | | |
|---|---|---|
| Birc2 | 1.054818 | 0.018978 |
| Egr2 | 4.890629 | 0.019792 |
| C330011M18Rik | 3.062187 | 0.0205 |
| Abhd14a | 3.228564 | 0.021022 |
| Trav14n-3 | 4.776638 | 0.021022 |
| Ggct | 1.201207 | 0.021022 |
| Egr1 | 2.031614 | 0.021416 |
| Mcoln2 | 2.041385 | 0.022436 |
| 9130019O22Rik | 2.691536 | 0.022549 |
| Gm17173 | 5.158085 | 0.022879 |
| Klrd1 | 1.319259 | 0.023979 |
| Rnf144a | 3.170648 | 0.023994 |
| Slc2a9 | 1.637185 | 0.023994 |
| Sik1 | 2.309241 | 0.025315 |
| Casp3 | 1.116233 | 0.02536 |
| Zfp7 | 1.486469 | 0.025889 |
| Gm11775 | 2.615296 | 0.026245 |
| Gm49004 | 1.893869 | 0.026245 |
| Fes | 1.35603 | 0.026759 |
| Gas2l3 | 2.790885 | 0.027359 |
| Gm16675 | 3.018773 | 0.027845 |
| Ubtd2 | 3.271827 | 0.028088 |
| Zfp948 | 3.78703 | 0.028876 |
| Cyth3 | 1.06176 | 0.030383 |
| Ntf5 | 3.50029 | 0.030383 |
| Sft2d2 | 1.095261 | 0.030458 |
| Lig4 | 2.710782 | 0.030458 |
| Cenpa | 2.588078 | 0.030459 |
| Gm44641 | 3.09145 | 0.030722 |
| AW549877 | 1.320012 | 0.031428 |
| Acoxl | 1.092486 | 0.031582 |
| Gm43042 | 1.125295 | 0.032614 |
| Gm17056 | 4.447523 | 0.033808 |
| Pdp1 | 1.637505 | 0.036477 |
| 4930505O20Rik | 4.560369 | 0.036717 |
| Rasgef1c | 2.089432 | 0.036717 |
| Rps26-ps1 | 1.741871 | 0.036976 |
| Tyrobp | 1.087746 | 0.037073 |
| Fam174b | 1.677018 | 0.037073 |
| Gm11767 | 1.972068 | 0.037971 |
| Cblb | 1.202986 | 0.038716 |
| Spo11 | 2.715193 | 0.040457 |
| 2700062C07Rik | 1.304598 | 0.040457 |
| Gm47571 | 2.421928 | 0.040457 |
| Itgav | 1.232708 | 0.040559 |
| Gm48430 | 1.95179 | 0.041389 |
| Trbv31 | 4.709295 | 0.041487 |
| Matk | 1.887288 | 0.042519 |
| Ccdc112 | 2.424079 | 0.0427 |
| Gm49359 | 1.577839 | 0.043574 |
| Nr4a2 | 2.071877 | 0.043789 |
| Ckm | 2.559892 | 0.044383 |
| Rasgef1a | 1.9702 | 0.04661 |
| Rtn4rl1 | 1.482473 | 0.046831 |
| Cd200 | 1.696852 | 0.049049 |
| Xaf1 | 1.549737 | 0.049049 |
| Rbm11 | 1.773925 | 0.049794 |
