## Supplementary material for "T cell receptor is required for differentiation but not maintenance of intestinal intraepithelial lymphocytes": Table S5

### Table S5. Differentially expressed genes between E8$_{I}^{WT(\textit{Trac})}$ and E8$_{I}^{\Delta(\textit{Trac})}$ nIELs

| Overexpressed in E8$_{I}^{WT(Trac)}$ | | | Overexpressed in E8$_{I}^{\Delta(Trac)}$ | | |
|---|---|---|---|---|---|
| **gene** | **ave_logFC** | **p_val_adj** | **gene** | **ave_logFC** | **p_val_adj** |
| Trac | 1.651493 | 2.71E-26 | Trdc | -4.53865 | 1.26254E-46 |
| Spry2 | 2.094503 | 1.23E-21 | Cish | -1.27063 | 7.03338E-10 |
| Fcrl1 | 8.550116 | 2.84E-16 | A130014A01Rik | -4.93386 | 1.30022E-09 |
| Gm1821 | 1.164367 | 6.83E-11 | Bbs10 | -7.48837 | 2.01595E-09 |
| Tigit | 1.679362 | 7.03E-10 | Plac8 | -1.16391 | 3.38171E-08 |
| Cd200r2 | 1.165216 | 4.22E-09 | Thy1 | -2.31287 | 4.3433E-08 |
| Klhdc9 | 7.42437 | 4.22E-09 | Syk | -6.91316 | 1.66875E-07 |
| Rpl36-ps2 | 6.328376 | 4.21E-08 | Lat | -1.18016 | 1.93934E-07 |
| Cela1 | 1.21494 | 4.54E-08 | Krt18 | -8.56492 | 3.91255E-07 |
| Smim10l2a | 4.245129 | 5.05E-08 | Scin | -1.94765 | 9.48968E-07 |
| Klrb1a | 1.827496 | 1.67E-07 | Gm47322 | -6.45336 | 2.60109E-06 |
| Gm28112 | 1.008876 | 1.93E-07 | Kbtbd13 | -6.45716 | 2.99673E-06 |
| Bcl2a1d | 1.643027 | 1.96E-07 | 5031439G07Rik | -4.1797 | 3.08547E-06 |
| Ikzf2 | 1.072478 | 2.2E-07 | Cd63 | -3.00503 | 3.08547E-06 |
| Nr4a2 | 2.143791 | 2.72E-07 | Gm42872 | -5.24863 | 5.84823E-06 |
| Cntln | 2.19592 | 3.91E-07 | Gm39091 | -6.42539 | 1.77748E-05 |
| Cnih3 | 4.192692 | 3.91E-07 | Thnsl2 | -4.97354 | 3.23055E-05 |
| Sh2d2a | 1.183584 | 8.47E-07 | B9d1 | -6.32686 | 3.26977E-05 |
| Arhgap11a | 4.707478 | 1.62E-06 | Cdc14b | -3.06993 | 3.67793E-05 |
| Apol10a | 6.531363 | 1.88E-06 | Gzmk | -1.00806 | 8.87897E-05 |
| Gm156 | 2.721936 | 2.24E-06 | Susd2 | -5.59168 | 0.000233757 |
| Bcl2a1b | 1.163319 | 2.37E-06 | Gm26353 | -6.01318 | 0.000274643 |
| Bcl2l15 | 1.619873 | 3E-06 | Pdzk1 | -7.3396 | 0.000320842 |
| Gjb2 | 4.760034 | 3.73E-06 | Rnf146 | -1.10054 | 0.000595956 |
| Gm17928 | 1.431437 | 3.73E-06 | Tpcn1 | -2.57531 | 0.000658268 |
| Sft2d2 | 1.474215 | 4.37E-06 | Gm30934 | -5.83734 | 0.000711777 |
| Pik3ap1 | 1.316334 | 5.85E-06 | Ifi208 | -1.46736 | 0.000723897 |
| Tex2 | 1.209198 | 5.98E-06 | Zscan30 | -5.80015 | 0.000810335 |
| Trav6-1 | 8.050649 | 1.4E-05 | Atp8a2 | -1.8044 | 0.001098701 |
| Klhdc2 | 1.038815 | 1.76E-05 | Osm | -6.16468 | 0.001165619 |
| Ccrl2 | 1.647218 | 1.98E-05 | Gm45378 | -5.7467 | 0.001165619 |
| Hal | 3.449997 | 3.68E-05 | Fnbp1l | -3.07833 | 0.001165619 |
| Il2rb | 1.278441 | 7.47E-05 | Gm45867 | -2.1641 | 0.001168729 |
| Gm43042 | 1.033033 | 8.97E-05 | Galnt10 | -2.4545 | 0.001353819 |
| Map4k5 | 2.318255 | 9.73E-05 | Cd6 | -3.89704 | 0.001390277 |
| Ptpn6 | 1.312568 | 0.000137 | E430021H15Rik | -1.39668 | 0.001390277 |
| Pdgfb | 5.47043 | 0.000145 | Ctrl | -5.69373 | 0.001429243 |
| Igip | 2.471296 | 0.000256 | Gm8000 | -5.66916 | 0.001445218 |
| Trio | 1.65572 | 0.00026 | Acp2 | -1.75448 | 0.001881934 |
| Arhgef3 | 1.033652 | 0.000338 | Gm10509 | -5.3901 | 0.002097053 |
| Gm44937 | 3.799383 | 0.000372 | Cd226 | -1.89377 | 0.002118614 |
| Klra3 | 4.05604 | 0.000385 | Ptafr | -2.42438 | 0.002436268 |
| Dclre1a | 2.138831 | 0.000454 | Gm44745 | -4.37293 | 0.002618644 |
| Gm12474 | 4.680125 | 0.000472 | Mgat4a | -1.69174 | 0.002840022 |
| Crem | 2.195776 | 0.000486 | Bcl11b | -1.54578 | 0.002858117 |
| Gm38304 | 5.693536 | 0.000506 | Acss2 | -1.16265 | 0.003634139 |
| Usp46 | 1.656611 | 0.000541 | Vezf1 | -1.04997 | 0.004021531 |
| Nlrp1b | 5.537333 | 0.00055 | Trav15d-1-dv6d-1 | -6.06744 | 0.004101732 |
| Wls | 1.250482 | 0.000674 | Pdzd9 | -5.42141 | 0.004181892 |
| Mdfic | 1.145723 | 0.000728 | Cdo1 | -3.67976 | 0.005921496 |
| Arap3 | 1.213769 | 0.000821 | Trav15d-2-dv6d-2 | -4.01615 | 0.006417876 |
| Gm44175 | 1.071119 | 0.000992 | Trav15n-2 | -4.01615 | 0.006417876 |
| Gm37313 | 5.728253 | 0.001166 | Cxcr3 | -1.01362 | 0.006417876 |
| Gsap | 1.132776 | 0.001184 | Gm11869 | -5.30471 | 0.00755292 |
| Gm47571 | 2.698057 | 0.001462 | AA914427 | -5.07344 | 0.00855734 |
| Gm4032 | 2.935027 | 0.001462 | Gm19331 | -4.02333 | 0.009319369 |
| Arhgef25 | 2.190686 | 0.001809 | Fbxl21 | -3.59406 | 0.009494792 |
| A930002I21Rik | 6.382722 | 0.001971 | Gm43566 | -5.2179 | 0.011211491 |
| Ttc30a1 | 6.701831 | 0.002094 | Gbp11 | -5.19167 | 0.011374094 |
| Trav9d-3 | 2.018748 | 0.002097 | Hist1h4n | -4.66668 | 0.01169224 |
| Trav9n-3 | 2.018748 | 0.002097 | Eepd1 | -5.03142 | 0.011742266 |
| Pde7b | 1.861599 | 0.002375 | Fbxo8 | -1.10453 | 0.011742266 |

| | | | | | |
|---|---|---|---|---|---|
| Bend4 | 2.325691 | 0.002433 | Lncbate6 | -1.84667 | 0.012738966 |
| Tsga10 | 1.196677 | 0.002619 | Gm12713 | -5.14117 | 0.013551604 |
| Cables1 | 1.073842 | 0.002781 | Klf3 | -2.28268 | 0.013568717 |
| Itih5 | 1.691067 | 0.002781 | Rgs4 | -2.68301 | 0.014253307 |
| Ckm | 1.563373 | 0.002835 | Cox7a1 | -2.99604 | 0.014653697 |
| Nr4a1 | 2.516035 | 0.003536 | BE692007 | -2.24481 | 0.015664207 |
| Gm19557 | 1.169353 | 0.003572 | Gm48565 | -4.69275 | 0.01658852 |
| Kctd15 | 4.331551 | 0.00382 | Gm37352 | -3.38135 | 0.016781735 |
| Zmym6 | 1.595695 | 0.004102 | Cdkl4 | -5.81185 | 0.020335977 |
| Cd200 | 2.313767 | 0.006078 | Hist3h2ba | -2.16839 | 0.02065085 |
| Ace2 | 5.363982 | 0.006418 | Deptor | -2.11977 | 0.021642335 |
| Gm45716 | 1.398436 | 0.006627 | Mbtps2 | -1.19896 | 0.022154154 |
| Slc16a11 | 2.017222 | 0.008392 | Cd4 | -1.9196 | 0.023487106 |
| Gm10801 | 1.056987 | 0.008683 | A530041M06Rik | -1.52878 | 0.024581149 |
| Gm45640 | 3.758381 | 0.009084 | Gm49368 | -1.62591 | 0.025536346 |
| Cers6 | 1.427203 | 0.009356 | Gm42161 | -3.40853 | 0.026337231 |
| Naa16 | 1.142078 | 0.011211 | Gm10392 | -1.77524 | 0.026872804 |
| Zfx | 1.599695 | 0.011223 | Gmeb2 | -2.02454 | 0.030630494 |
| 4930431P03Rik | 2.610789 | 0.011223 | Emp3 | -1.06952 | 0.030630494 |
| Gm37124 | 3.526872 | 0.011541 | Inpp4b | -1.75433 | 0.031222696 |
| A530083M17Rik | 1.692958 | 0.011692 | Gm28791 | -1.02655 | 0.031222696 |
| Pidd1 | 2.467629 | 0.011742 | Mirt1 | -1.09175 | 0.031408479 |
| Gm45235 | 2.023999 | 0.012155 | Ifit1bl1 | -1.10279 | 0.032532433 |
| Gm17300 | 1.782229 | 0.01246 | Sit1 | -3.39506 | 0.034254991 |
| Asah2 | 3.111448 | 0.012703 | Uchl1os | -2.19954 | 0.034405101 |
| Plek | 1.905637 | 0.013253 | Zfp574 | -1.55884 | 0.041200011 |
| Gm43137 | 5.171314 | 0.013253 | Clec12a | -2.16814 | 0.042411778 |
| Lag3 | 1.051722 | 0.014253 | Ifitm10 | -1.27995 | 0.042678342 |
| Cyth3 | 1.065344 | 0.014253 | 1600012H06Rik | -1.32861 | 0.042843538 |
| Tspan4 | 2.155092 | 0.014253 | Muc1 | -4.83637 | 0.04396801 |
| Med14 | 1.352428 | 0.014598 | Trbj1-1 | -1.7668 | 0.046394178 |
| Trav16n | 2.107524 | 0.015334 | Gm42780 | -4.53285 | 0.048510052 |
| Crim1 | 2.693165 | 0.015334 | B4galnt2 | -1.38957 | 0.048510052 |
| Zfp595 | 1.562694 | 0.015742 | Fam53b | -1.65382 | 0.048924305 |
| Mettl9 | 1.161021 | 0.016583 | Trdc | -4.53865 | 1.26254E-46 |
| Cdkn2b | 4.452195 | 0.016758 | Cish | -1.27063 | 7.03338E-10 |
| Pom121 | 1.440646 | 0.017047 | A130014A01Rik | -4.93386 | 1.30022E-09 |
| Gm29112 | 3.285148 | 0.018774 | Bbs10 | -7.48837 | 2.01595E-09 |
| Gm43413 | 1.11705 | 0.018933 | Plac8 | -1.16391 | 3.38171E-08 |
| Pabpc4 | 1.906865 | 0.020651 | Thy1 | -2.31287 | 4.3433E-08 |
| Egr1 | 1.990024 | 0.020902 | Syk | -6.91316 | 1.66875E-07 |
| H60b | 1.952307 | 0.020941 | Lat | -1.18016 | 1.93934E-07 |
| Klrb1 | 1.701475 | 0.023204 | Krt18 | -8.56492 | 3.91255E-07 |
| Filip1 | 6.355295 | 0.023276 | Scin | -1.94765 | 9.48968E-07 |
| Kit | 1.196299 | 0.023361 | Gm47322 | -6.45336 | 2.60109E-06 |
| Gm11168 | 1.025009 | 0.023487 | Kbtbd13 | -6.45716 | 2.99673E-06 |
| 6030400A10Rik | 5.006698 | 0.023729 | 5031439G07Rik | -4.1797 | 3.08547E-06 |
| Pcdhb7 | 4.754488 | 0.025397 | Cd63 | -3.00503 | 3.08547E-06 |
| Ints14 | 1.582965 | 0.02603 | Gm42872 | -5.24863 | 5.84823E-06 |
| Itgav | 1.039502 | 0.02789 | Gm39091 | -6.42539 | 1.77748E-05 |
| Ccdc102a | 1.119039 | 0.029585 | Thnsl2 | -4.97354 | 3.23055E-05 |
| Usp38 | 1.29133 | 0.029926 | B9d1 | -6.32686 | 3.26977E-05 |
| Pheta2 | 1.103337 | 0.030221 | Cdc14b | -3.06993 | 3.67793E-05 |
| Cd40lg | 4.582232 | 0.032083 | Gzmk | -1.00806 | 8.87897E-05 |
| Gm33104 | 1.142824 | 0.033322 | Susd2 | -5.59168 | 0.000233757 |
| Klra1 | 1.382559 | 0.033322 | Gm26353 | -6.01318 | 0.000274643 |
| Hpf1 | 1.157305 | 0.034255 | Pdzk1 | -7.3396 | 0.000320842 |
| Spred1 | 2.414797 | 0.034741 | Rnf146 | -1.10054 | 0.000595956 |
| Gm3650 | 3.878987 | 0.035173 | Tpcn1 | -2.57531 | 0.000658268 |
| Fam92a | 1.220107 | 0.037203 | Gm30934 | -5.83734 | 0.000711777 |
| Kbtbd7 | 1.920862 | 0.037684 | Ifi208 | -1.46736 | 0.000723897 |
| Naprt | 1.655793 | 0.0412 | Zscan30 | -5.80015 | 0.000810335 |
| Gm44037 | 2.876672 | 0.041263 | Atp8a2 | -1.8044 | 0.001098701 |
| Gm45894 | 3.898248 | 0.041696 | Osm | -6.16468 | 0.001165619 |
| Adh1 | 1.836137 | 0.041785 | Gm45378 | -5.7467 | 0.001165619 |
| Dclk2 | 2.886998 | 0.043968 | Fnbp1l | -3.07833 | 0.001165619 |
| Zfp65 | 1.799708 | 0.044198 | Gm45867 | -2.1641 | 0.001168729 |

| | | | | | |
|---|---|---|---|---|---|
| Gm20707 | 1.776769 | 0.044671 | Galnt10 | -2.4545 | 0.001353819 |
| Gm10717 | 1.016993 | 0.044708 | Cd6 | -3.89704 | 0.001390277 |
| Crybg1 | 1.237992 | 0.044708 | E430021H15Rik | -1.39668 | 0.001390277 |
| Tmem186 | 1.184771 | 0.046619 | Ctrl | -5.69373 | 0.001429243 |
| 1700037H04Rik | 1.390203 | 0.046891 | Gm8000 | -5.66916 | 0.001445218 |
| St7l | 1.267517 | 0.047474 | Acp2 | -1.75448 | 0.001881934 |
| Tssk6 | 3.311044 | 0.047474 | | | |
| Zfp850 | 2.04788 | 0.04851 | | | |
| Edem3 | 1.512383 | 0.049055 | | | |
| Gm10720 | 1.092274 | 0.049493 | | | |
| Slc35f5 | 1.934453 | 0.049793 | | | |
